# Pangenomic context reveals the extent of intraspecific plant NLR evolution

**DOI:** 10.1101/2024.09.02.610789

**Authors:** Luisa C. Teasdale, Kevin D. Murray, Max Collenberg, Adrian Contreras-Garrido, Theresa Schlegel, Leon van Ess, Justina Jüttner, Christa Lanz, Oliver Deusch, Joffrey Fitz, Regina Mencia, Rosanne van Velthoven, Hajk-Georg Drost, Detlef Weigel, Gautam Shirsekar

## Abstract

Nucleotide-binding leucine-rich repeat (NLR) proteins are a major component of the plant immune system, which directly or indirectly detect molecular signals of pathogen invasion. Despite their critical role, the processes by which NLR genes diversify remain poorly characterised due to the extraordinary sequence, structural, and regulatory variability of NLRs, even among closely related individuals. To understand the evolution of NLR diversity in *Arabidopsis thaliana*, we leverage graph-based methods to define pangenomic NLR neighbourhoods in 17 genetically diverse genomes. We integrate full-length transcript and transposable element information to exhaustively annotate all intact and degraded NLRs, enabling exploration of the processes that underpin the birth, death and maintenance of NLR diversity within a species. Our main finding is that many uncorrelated mutational processes create NLR diversity, and that there is no single metric that captures on its own the true extent of NLR structural and sequence variation. This immense diversity in plant immune system diversification allows populations to survive the constant onslaught of pathogens, not unlike vertebrate adaptive immunity, where variation is also generated by a variety of complementary mechanisms, albeit at the level of individuals.

## Introduction

All organisms have to defend themselves against a multitude of enemies, for which they use both physical and biochemical means. The latter often rely on detecting alien molecular signals that initiate countermeasures by the attacked host. While plants lack an adaptive immune system in the vertebrate sense, they benefit from extensive population-level genetic diversity of immune genes (Brown and Tellier 2011), with networks of interacting sensors and downstream factors that detect infection and trigger defence (Dodds and Rathjen 2010; Jones et al. 2024). As a result, components of the plant immune system both evolve more rapidly than the genome at large, but also maintain diversity for much longer than the rest of the genome, in co-evolution with multiple pathogens whose abundance and diversity fluctuate over space and time (Thrall and Burdon 2003; Thompson 2005; Koenig et al. 2019). Because of a general trade-off between immunity and plant vigour, there are also inherent limits to the number and diversity of immune genes (Karasov et al. 2017). Therefore, selection on immune genes can be both strong and ephemeral at the same time, with successive waves of diversification and selection maintaining an evolutionary balance both between plants and their pathogens, and between growth, defence, and avoidance of autoimmunity.

A major group of immune receptors are the NLR proteins (nucleotide-binding leucine-rich repeat proteins; alternatively NOD-Like Receptors), which detect effector proteins that help pathogens to evade or co-opt the primary, generalised stage of pattern-triggered immunity (Jones et al. 2024). The diversity of NLRs appears to match the enormous diversity of pathogen effector repertoires (Michelmore and Meyers 1998; Bakker et al. 2006; Clark et al. 2007). NLR genes vary in their primary domain composition, with some active NLRs lacking several canonical NLR domains, and they can be found in complex and dynamic gene clusters or occur in stable paired or single gene configurations. Finally, not all NLRs are directly involved in pathogen recognition, and some function as executor of helper NLRs downstream of sensor NLRs (Barragan and Weigel 2021).

Early analyses noted high nucleotide-level diversity of individual NLRs (Bakker et al. 2006) and highlighted the birth, divergence, and death of genes within clusters as potential key mechanisms underlying NLR evolution (Michelmore and Meyers 1998). These patterns have been confirmed at increasing scales in the decades since (Clark et al. 2007; Van de Weyer et al. 2019; Lee and Chae 2020; Prigozhin and Krasileva 2021), including recent insights into epigenetic and regulatory variation (Tsuchiya and Eulgem 2013; Kawakatsu et al. 2016; Brabham et al. 2024; Sutherland et al. 2024). What remains less clear are the specific mechanisms by which plants generate and maintain functionally relevant NLR diversity (Barragan and Weigel 2021).

While NLRs often come to the fore when one looks for a high density of structural variants that disrupt synteny between closely related genomes (Jiao and Schneeberger 2020), one can understand the evolution of the entire NLR family only by considering both population diversity and the genomic context of all NLR genes. Here, based on evidence-based rigorous annotation and epigenetic profiles of the individual genomes, we characterise the NLR gene complement in genomic and population context across a diverse collection of 17 genomes that represent range-wide haplotype diversity of *A. thaliana*. We used a novel pangenome graph-based principled approach for delineating the complex genomic regions surrounding NLRs, what we call “NLR neighbourhoods”. Importantly, our methods are robust to the rampant structural variation typical for many regions containing multiple NLRs. Using our pangenomic neighbourhood approach, along with high-confidence annotations of NLRs, pseudogenes, and transposable elements, a much richer picture of immune system evolution emerges. Notably, NLR diversification and evolution defy classification by simple measures or metrics, but can only be understood with a holistic view.

## Results

### Defining the pangenomic context of NLRs

To characterise the dynamics of NLR evolution in *A. thaliana,* we selected 17 accessions to represent the genetic diversity across the Western Palearctic, based on geographic stratification and previously identified haplotype sharing groups (Shirsekar et al. 2021) (Fig. 1a). We assembled and scaffolded contigs from PacBio HiFi reads into chromosomes with the aid of a BioNano optical map for one of the accessions (at9852). We annotated protein coding genes using both homology and full-length cDNA expression evidence from PacBio Iso-Seq data, and annotated transposable elements (TEs) using both homology and curation of novel repeat families. We annotated up to 29,321 protein coding genes per accession (including pseudogenes but not including TE protein genes), with an average of 35.1 Mb of repeats, and a combined total of 3,789 NLR genes (Sup. Fig. 1). As expected, all genomes show a high degree of global synteny (Sup. Fig. 2) (Lian et al. 2024). We took an integrated approach to the annotation of the NLR genes, combining multiple sources of evidence including Iso-Seq data with manual curation to produce high-confidence NLR gene models (see Methods and Sup. Fig. 3 for full details). This annotation approach improves upon previous work that relied on single-accession references, target capture techniques, and short-read RNAseq. We also manually annotated and verified NLR pseudogenes and partial NLR copies that could not be annotated by automated approaches, many of which likely have compromised activity, allowing us to examine mutational processes that inactivate NLRs.

**Figure 1:**
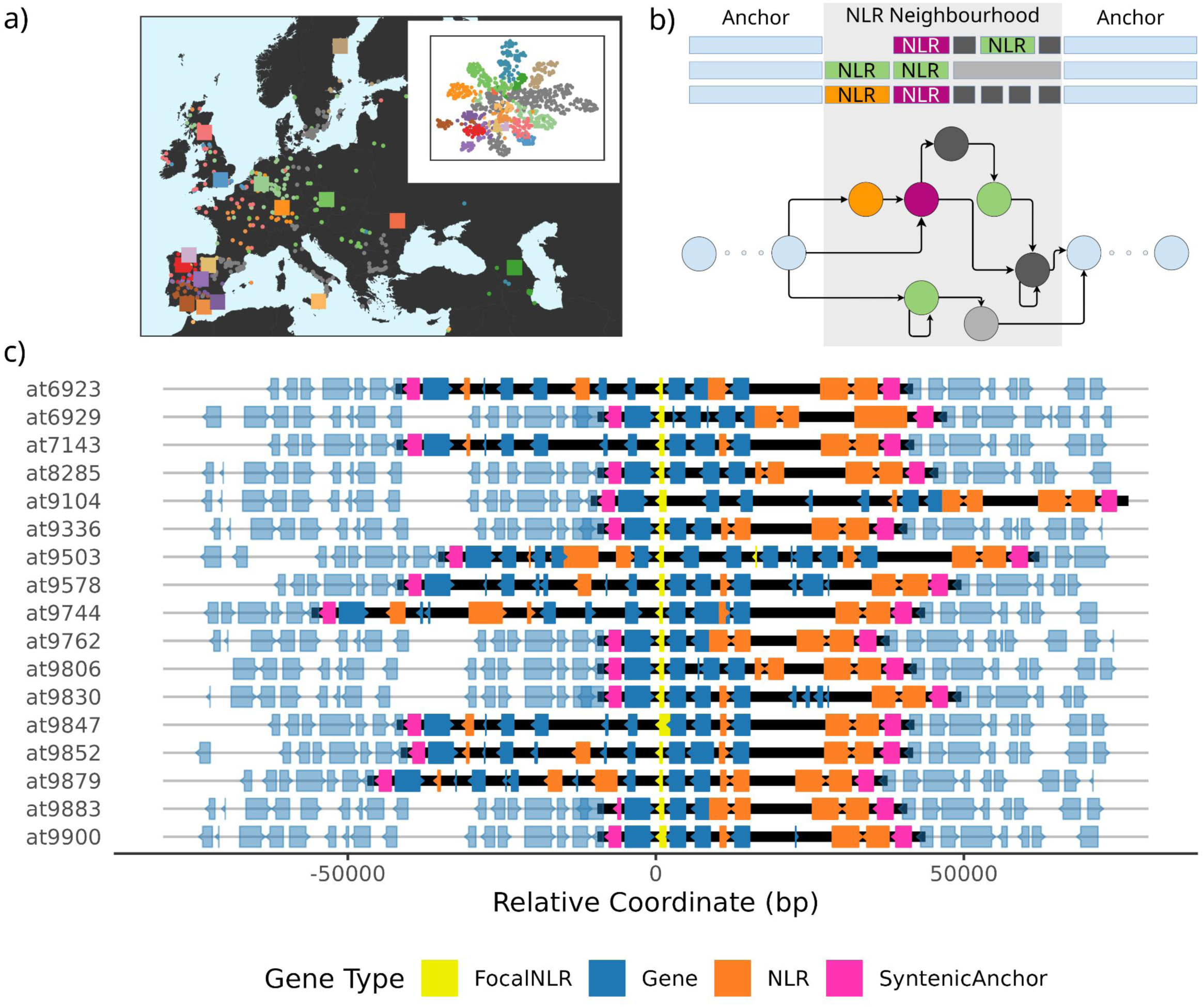
Defining NLR neighbourhoods across 17 *Arabidopsis thaliana* genomes. **a,** Our 17 accessions (squares) are broadly representative of both the geographic and population genetic distributions of the broader 1001 Genomes collection (circles; Alonso-Blanco et al. 2016); inset shows the UMAP embedding of 17 accessions in the co-ancestry space of all 1135 accessions (adapted from Shirsekar et al 2020). Colours in a) represent the haplotype sharing group from Shirsekar et al (2020). **b,** Universal, pangenome-wide syntenic anchors define NLR neighbourhoods. **c,** Representative example of an NLR neighbourhood, highlighting accurate delineation of the borders of NLR-containing variable regions, regardless of structural or presence/absence variation. Black bars indicate pangenomic neighbourhoods, syntenic anchors in pink. A fixed-sized window approach centered on a focal NLR (in yellow) would produce a very different outcome.

Given that TEs and the epigenetic landscape can influence NLR function (Tsuchiya and Eulgem 2013; Sutherland et al. 2024), we aimed to take a principled approach to taking the genomic environment of NLRs into account. To do so, we devised a method for consistent delineation of NLR regions that is robust to frequent presence-absence variation. Using a pangenomic strategy we defined “NLR neighbourhoods” as regions that contain at least one NLR in at least one accession, bordered by pangenome-wide syntenic anchors (Fig. 1b, c). These regions vary in complexity: in the simplest cases they consist of a single syntenic NLR, but they can also be very large and among the most complex genic portions of the pangenome. Compared to defining NLR clusters as a region that includes sequences up and downstream of an NLR or clustering NLRs based solely on genomic proximity (Holub 2001; Lee and Chae 2020; Prigozhin and Krasileva 2021), our approach guarantees that all structural variation surrounding NLRs is encompassed within a specific neighbourhood, even in cases where NLRs are entirely absent in some accessions, facilitating study of complete losses as well as *de novo* emergence of NLR loci.

Across all genomes, we identified 121 NLR neighbourhoods. These generally are similar to other euchromatic regions with respect to protein-coding gene density and steady-state cytosine methylation (Sup. Fig. 4). However, NLR neighbourhoods tend to have a higher TE density, with on average younger TEs as inferred from divergence from the family consensus (Sup. Fig. 4).

### NLR neighbourhood complexity and variability vary greatly

NLR neighbourhoods differ greatly in average length, from 1.9 kb to over 900 kb, although most are smaller than 50 kb (Fig. 2a). Similarly, while the length of some neighbourhoods varies considerably between accessions, this is not the case for most (Fig. 2a). The neighbourhoods are widely variable in NLR content, ranging from neighbourhoods with only NLR fragments present in just a few accessions, to neighbourhoods with several NLRs that themselves differ greatly in size and sequence across accessions (Fig. 2b). A numerous class, about 33%, comprises single-copy NLRs with highly conserved synteny.

**Figure 2:**
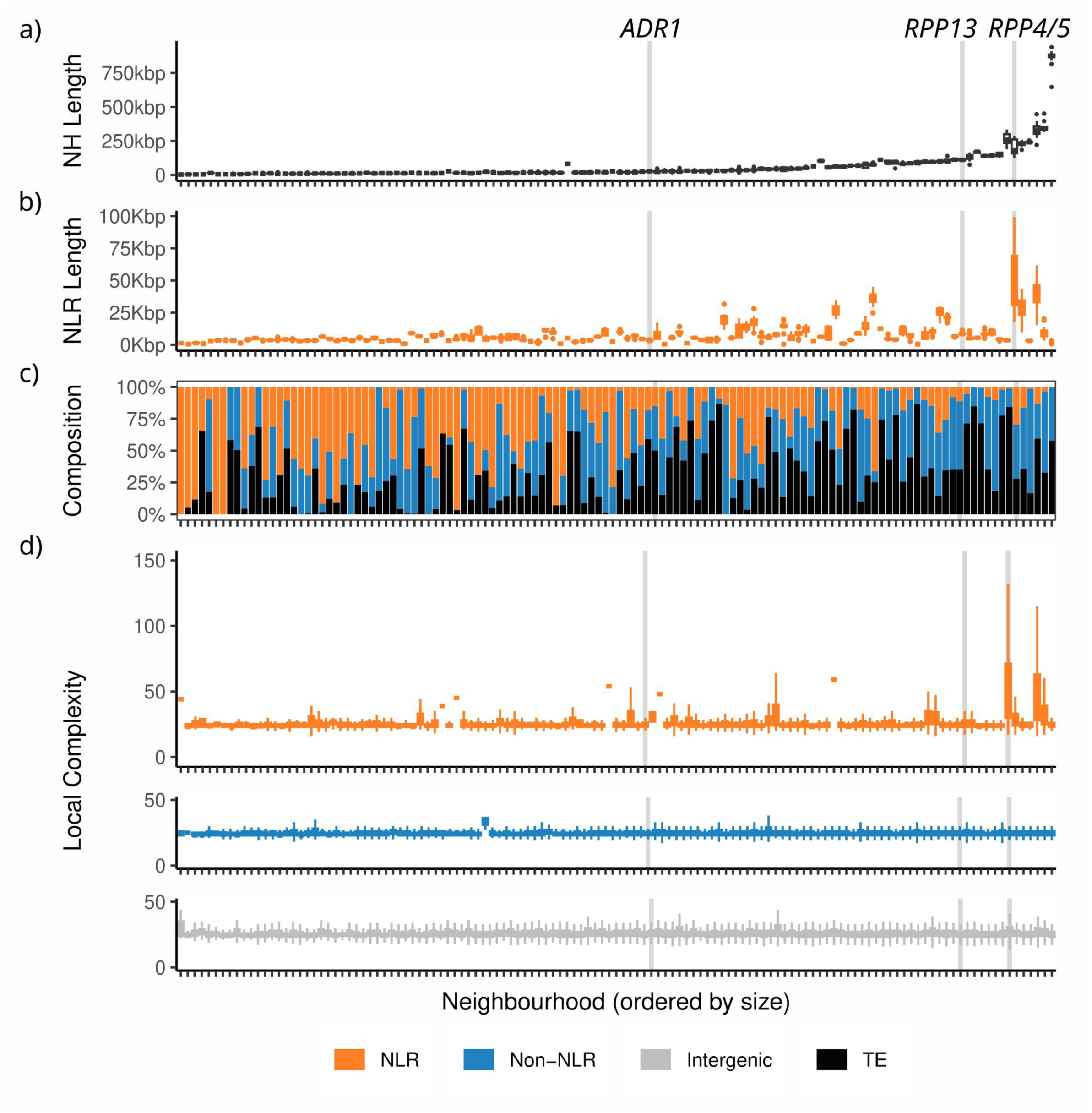
NLR neighbourhoods across 17 genomes. **a-c,** NLR neighbourhoods vary across accessions in their overall length (a), combined length of NLR sequences (b), and the composition of their annotated space (c). **d,** Pangenome local complexity varies within and between neighbourhoods, and where elevated, is predominantly elevated in NLR genes themselves rather than non-NLR genes or intergenic space. In all subplots, neighbourhoods are ordered by their mean length across accessions.

There is also substantial variation in the relative proportions of NLRs, non-NLR protein coding genes, and TEs making up NLR neighbourhoods (Fig. 2b, c). Notably, there is no simple relationship between the size of a neighbourhood and the number of NLRs it contains. While larger neighbourhoods on average have a higher fraction of TEs, potentially indicating that these represent a genomic context that is more tolerant to TE activity, many counterexamples exist (Fig. 2c). The helper NLRs of the *ADR1* and *NRG* families as well as loci conferring resistance to generalist bacterial pathogens such as *RPM1*, *RPS5*, or *RPS2*, on average occur in smaller, less variable neighbourhoods. The larger, more variable neighbourhoods typically contain NLR genes that are co-evolving with highly specialised oomycete pathogens, such as *RPP1* and *RPP4/5*, which encode resistance to *Hyaloperonospora arabidopsidis*, but again, with notable exceptions: the neighbourhood (chr2_nbh01) with the highest average length is defined by two NLR fragments in a sea of highly complex TE insertions.

### Pangenomic complexity in NLR neighbourhoods is centred on NLRs

The degree of variation and structural complexity varies not only among NLR neighbourhoods, but also within them. We quantified the local complexity within neighbourhoods using a pangenome graph metric that summarises the degree of structural variation around a sequence (“node”) across all occurrences in the pangenome, what we call “node radius” (see Sup. Fig. 5). This metric represents the local connectedness of sequences within a pangenome graph. It increases especially with rearrangements, duplication, and translocations, as these increase the number of other sequences that can be reached from the focal node. In neighbourhoods with increased local complexity, this diversity is often centred on the NLRs (Fig. 2d; Fig 3). This observation suggests that rather than diversity being a consequence of the surrounding genomic context– for example, because of DNA sequences that are particularly prone to double-strand breakage or attracting TE insertions– it is often driven by duplication and rearrangement of the NLRs themselves. These patterns are exemplified by the *RPP4/5* locus, where a high degree of copy-number, domain, and single-nucleotide variation greatly increases our complexity metric over most of the numerous NLRs in this neighbourhood, while TEs, including recent insertions, contribute less to pangenomic complexity (Fig. 3). In contrast, in the *RPP13* neighbourhood, conserved gene structure and limited copy-number variation only slightly increases local complexity, despite sequences encoding the LRR domains of the NLRs varying greatly (Sup. Fig. 6).

**Figure 3:**
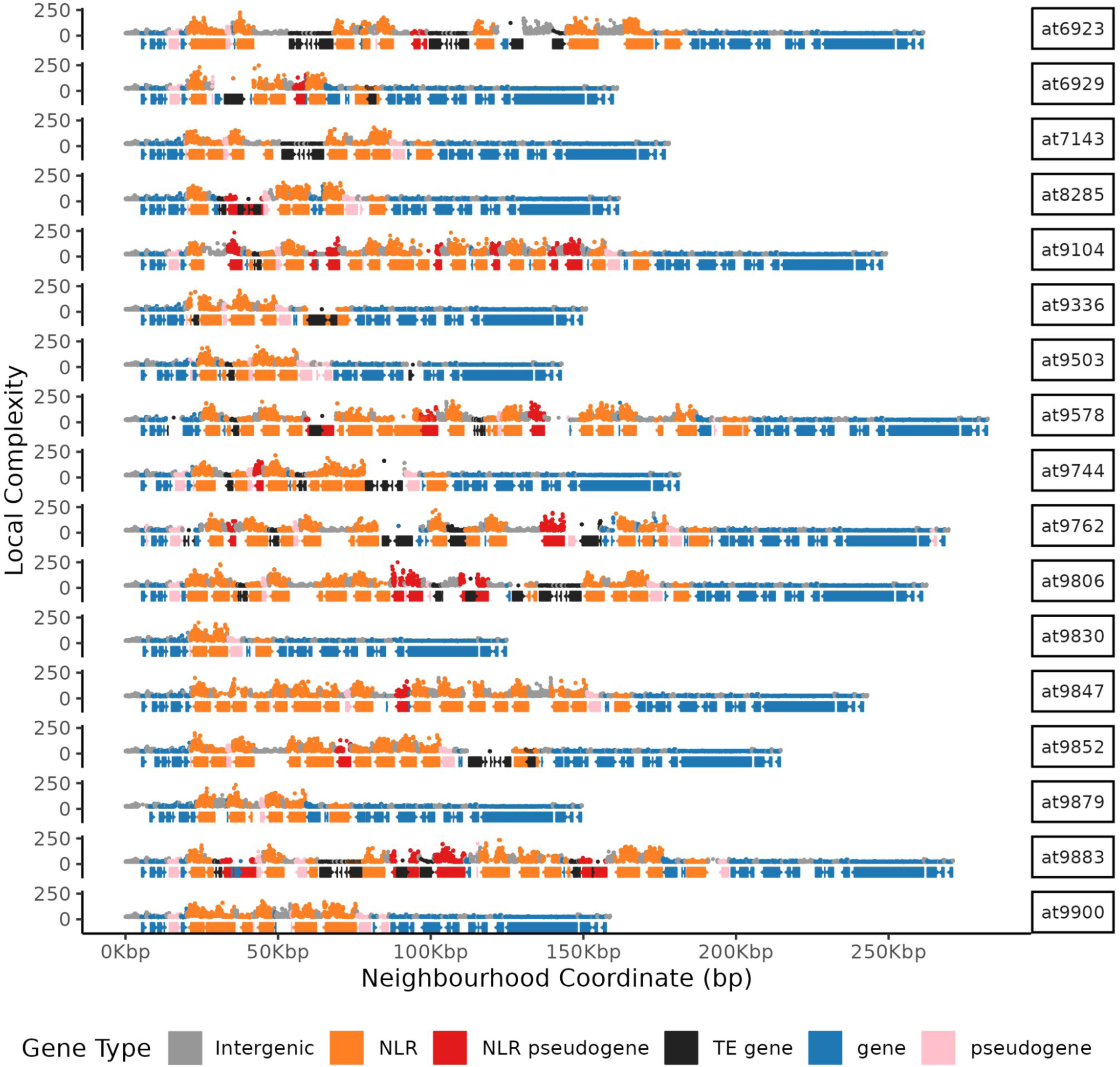
Pangenome complexity of the *RPP4/RPP5* NLR neighbourhood. The neighbourhood containing RPP4/RPP5 is among the most complex, and the highly elevated local complexity is focused on NLR genes and pseudogenes. Each per-accession track consists of both a trace of local pangenomic complexity (above the abscissae), and a representation of the gene annotation (below the abscissae). Individuals show extreme haplotypic diversity of NLRs, which is reflected in highly elevated local complexity focused on NLR genes and their immediate surrounds, while TE genes or other protein coding genes show little elevation in complexity above the genome background.

### NLRs are diverse in how they are diverse

That local complexity in NLR neighbourhoods is often centred on the NLRs themselves is consistent with rampant structural variation in many NLR genes. However, structural variation is only one of many forms of diversity. We used several complementary metrics to measure other axes of NLR diversification, such as pairwise nucleotide distance, population-wide frequency of high-impact mutations, isoform diversity, Shannon entropy of amino acid variation, diversity of NLR-associated domains (Sup. Fig. 17), and copy number variation (see Methods). These metrics were calculated for homologous clusters of NLR sequences (see Methods). Across these metrics, NLR gene families differ in both their main axes and degrees of diversity (Fig. 4a, b). Importantly, no single metric captures the majority of NLR diversity on its own (Sup. Fig. 7). This also applies to NLRs demonstrated to confer specific disease resistance, although they are among the more diverse NLRs by most metrics. For example, *RPP13* is highly diverse in sequence but not isoforms or copy number, whereas *DAR4* shows the opposite pattern, while *RPP4* is consistently extreme in nearly all metrics (Sup. Fig. 14).

**Figure 4:**
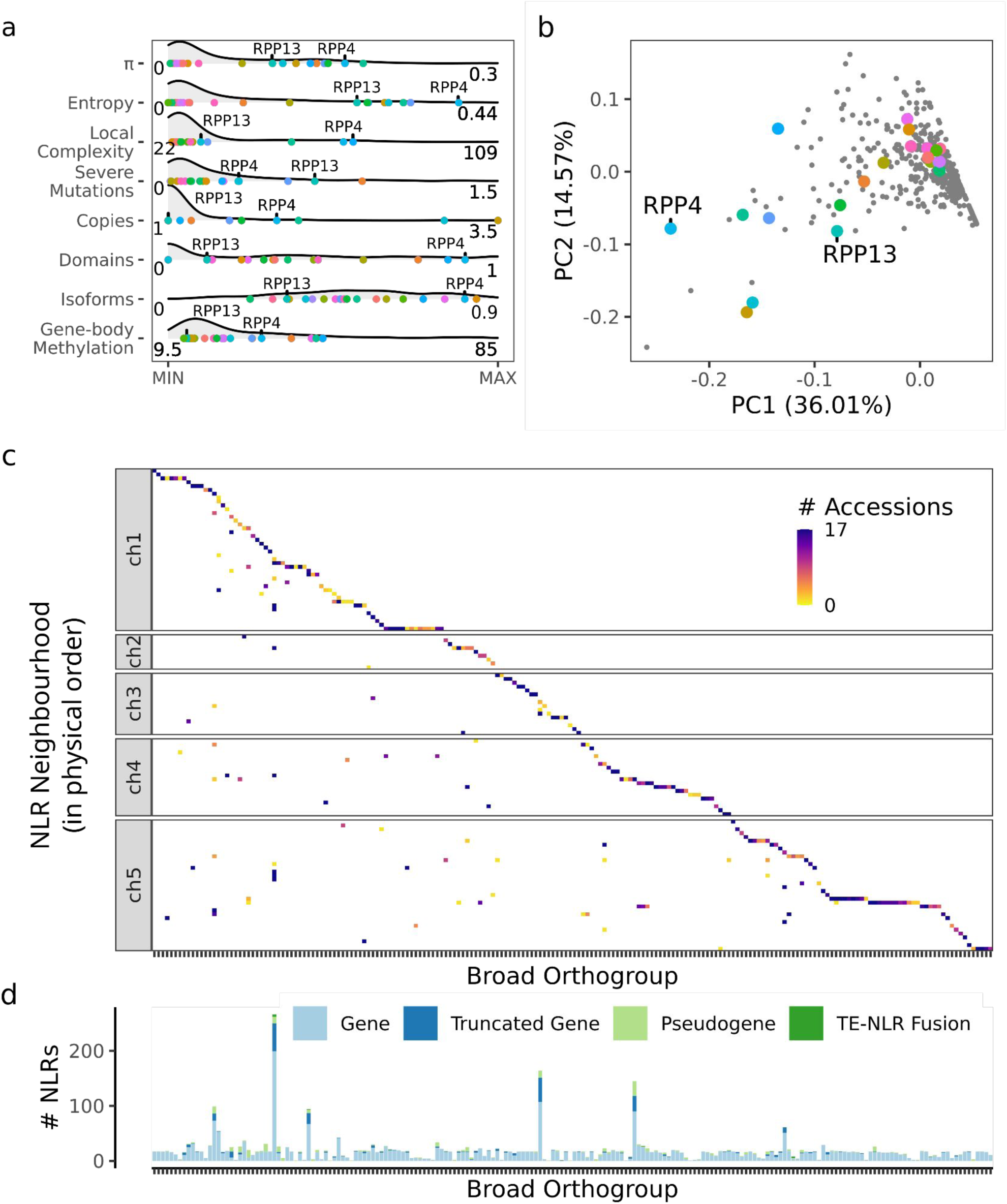
Linking genomic context to gene families. **a,** Variation of OG70 diversity across different axes. If not stated otherwise, always for all OG70 members across all accessions (a). π, mean nucleotide pairwise distance; Entropy, mean Shannon entropy of column states within an amino acid multiple sequence alignment, excluding positions with fewer than 10 non-gap characters; Local Complexity, mean pangenome local complexity; Severe Mutations, sum of allele frequencies of mutations with severe predicted consequences among the 1001 Genomes collection for OG70s present in at9852; Copies, mean number of copies per accession; Domains, mean Simpson’s index of diversity of NLR-associated domains detected by NLRtracker; Isoforms, mean of Simpson’s index of diversity of isoform variation affecting the open reading frame of each transcript; Gene-body Methylation, mean percentage of methylated CpG sites. **b,** PCA of OG70 diversity across each of the axes from (a) highlights that diversity varies in both degree and class between orthogroups. In both (a) and (b), coloured dots represent NLR genes previously identified as encoding resistance to a specific pathogen (see Sup. Table 2), which on the whole are distributed throughout the range of each diversity metric (a), and are relatively evenly distributed throughout the multivariate diversity space (b). **c,** The distribution of broad OGs across the 121 NLR neighbourhoods, ordered by chromosome positions (y-axis), and the first occurrence of the OG (x-axis). **d,** Total gene count per OG across accessions, classified by pseudogenisation state, ordered as in (c).

Transcript isoform variation is an under-appreciated axis of NLR diversity. It is only weakly correlated with domain diversity (adj. R^2^ 0.10, p=1.1e-9) and gene length (adj R^2^ 0.18, p=1.3e-15) (Sup. Fig. 8). Instead, it is a specific property of individual NLR gene families (adj. R^2^ 0.45, p<2.2e-16) (Sup. Fig. 9a). Therefore, genes that are highly conserved based on genomic DNA-based metrics can still produce diverse proteins across accessions through the expression of alternative transcripts (eg. the bacterial effector AvrRPS4 recognizing NLR gene *Rps4*). NLRs feature more isoform variation than non-NLR genes in the same neighbourhood, indicating that this increased diversity is specific to NLRs rather than epigenetic cues affecting a broader genomic context (Sup. Fig. 10).

However, we did find that methylation in NLR neighbourhoods is driven by TE presence rather than methylation of the NLRs themselves (Sup. Fig. 22).

### New insights into how NLR diversity has been generated

Turning to the processes and possible molecular mechanisms generating NLR diversity, we first looked at the gain and loss of NLR genes. We initially defined 204 more broadly related NLRs orthogroups (broad OGs). A little under half (88) were split into more finely delineated groups with at least 70% protein sequence identity (OG70s), for a total of 371 OG70s (Sup. Fig. 18). Some NLR neighbourhoods with multiple OGs are likely evolutionarily old, as there are typically only single members of multiple OGs. In other cases, such as the *RPP1* and *RPP4/5* neighbourhoods, copy number expansions have likely occurred after speciation, and these neighbourhoods generally contain few broad OGs, but with each being represented by multiple copies, indicative of an ongoing and active copy number expansion process (Fig. 4c). To be more specific, many NLR neighbourhoods (45%) contain multiple broad OGs, but most broad OGs, 72%, are found in only one neighbourhood (Fig. 4c).

In terms of conservation of individual OGs and OG70s, fewer than half (47%) of broad OGs, and an even smaller fraction of OG70s (18%), are present in all accessions. Conversely, OG70s are more likely to be restricted to a single accession (18% of OG70s) than broad OGs (4%) (Sup. Fig. 18). Some relatively rare OG70s represent long-range translocations of NLRs into new, distant neighbourhoods, and are likely recent and TE-mediated (Fig. 4c; Sup. Fig. 16).

A special case is represented by neighbourhoods with paired, yet highly divergent OGs that encode proteins forming obligate heteromers (van Wersch and Li 2019). Complete duplication of such paired genes (Saucet et al. 2015) and different pairs of alleles at the *CHS3*-*CSA1* locus having different biochemical functions are indicative of functional linkage of paired NLR genes being under strong purifying selection. In agreement, tight genetic linkage between paired NLRs is maintained across accessions in our dataset. Finally, as in the *A. thaliana* reference genome, we did not find cases where CNLs and TNLs share a neighbourhood, something that has been observed in other species (Ameline-Torregrosa et al. 2007).

An NLR neighbourhood with clear examples of both NLR birth and death is the neighbourhood containing *ADR2*. This neighbourhood has multiple copies of a single broad OG, several of which have undergone independent duplication and pseudogenisation (Fig. 5 a,b). Two separate pseudogenisation events have occurred in this neighbourhood, once in OGbroad_cluster-5 and once in OGbroad_cluster-2. In both cases intact copies still exist in some accessions. No accession has the full complement of all six OG70s found in this neighbourhood across accessions.

**Figure 5:**
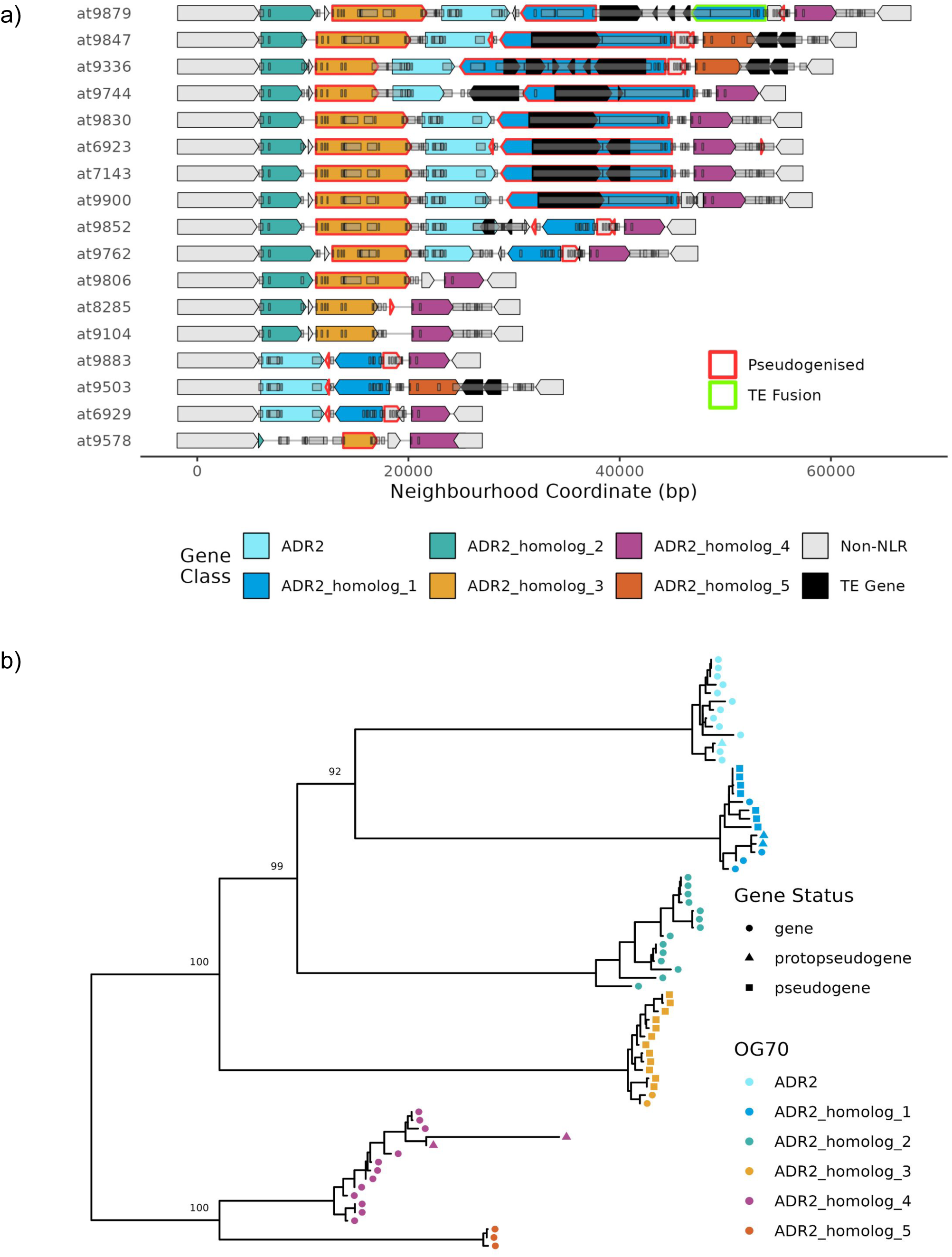
*ADR2* as an example of a neighbourhood with birth and death of NLRs. **a,** a local map of NLR genes in the neighbourhood in each accession, with genes coloured by OG70 membership. While some OG70s are intact in most accessions (ADR2, ADR2 Homolog 4), others show presence/absence variation (ADR2 Homologs 2, and 5), and/or frequent pseuodgenisation (ADR Homologs 3 and 1). Most accessions have 2-3 intact NLRs and 1-2 pseudogeneised NLRs, however some accessions have heavily reduced haplotypes containing no pseudogenes (e.g. at9578, at6929). **b,** A gene tree of NLRs in this neighbourhood, with annotated pseudogenisation events. The gene tree shows that there have been at least two separate pseudogenisation events within the neighbourhood within both ADR homologs 3 and 1.

Finally, there are examples of inversions and other structural variants generating novelty, such as at9879, where a TE is inserted into an NLR and the resulting new transcript encodes a fusion of the N-terminus of the NLR with a DUF1985 domain containing portion of a VANDAL2 TE (Fig. 5a).

### Pseudogenisation is frequent but pseudogenised alleles rarely persist or spread

Compared to other genes, a major feature that makes NLR genes stand out is the large fraction of genes with mutations that interrupt the open reading frame. Notably, genes that encode only some of the domains found in canonical NLRs can have biological function (Nishimura et al. 2017; Liang et al. 2019), making it much more difficult to decide if a truncated gene has potential for meaningful activity. Of all annotated NLR genes, 7.5% have a truncated open reading frame relative to allelic copies in other accessions, with single premature stop codons or frameshift mutations that are not fixed in the population. An additional 8.4% have multiple disabling mutations or produce transcripts that no longer encode any canonical NLR domain (Fig. 4d). Genes in the latter class are more likely to have only limited expression compared to the genes with single large-effect mutations (Sup. Fig. 19), consistent with these genes being further along the path to pseudogenisation. The causes of pseudogenisation include a variety of mutational processes, such as single-nucleotide variants and small indels causing in-frame stop codons, as well as, more rarely, TE insertions or larger deletions (Sup. Fig. 11). Single-copy conserved NLRs are less likely to be pseudogenised (Fig. 4d), suggesting that, as a group, they are more likely to be essential.

While NLRs seem to have a higher burden of severe mutations than non-NLR genes, individual deleterious variants are most often found in only a single accession. To more accurately estimate the population frequency of large-effect mutations, we called single-nucleotide variants from short-read data of 1,135 individuals from the 1001 Genomes Project (Alonso-Blanco et al. 2016) against one of our annotated accessions (at9852). The combined occurrence (the product of allele frequency and number of alleles) of variants with predicted severe consequences varies considerably among NLRs, with some genes with known roles in disease resistance, such as *RPP13*, having an increased burden of large-effect mutations (Fig. 4a). This burden is also higher for NLRs than for other protein-coding genes (Sup. Fig. 12), albeit it only marginally, and it is not strongly correlated (R^2^=0.065) with the overall sequence diversity within NLR OG70s (Sup. Fig. 7).

### NLR neighbourhoods are enriched for recent TE activity

TE sequences occur more frequently in NLR genes than in non-NLR genes. Genome-wide, just over half of intact NLR genes contain at least one TE sequence, while it is closer to a quarter for non-NLR genes. While, judging by their size, most TE sequences in NLR genes appear to be incomplete, the difference between NLR and non-NLR genes also holds when considering only intact TE insertions, with 3.9% of NLRs and 2.6% of non-NLR genes containing intact TEs, and when considering only TEs with clear homology to annotated TEs in Col-0 (NLRs: 33.1%, Non-NLRs: 14.5%). While in some cases, TEs within NLRs are conserved across accessions, in most cases the TEs within an NLR vary across accessions (Sup. Fig. 23).

NLR neighbourhoods are enriched for TEs, containing a wide variety of TE families. TEs within NLR neighbourhoods are on average younger than outside NLR neighbourhoods, as judged by formal models for dating intact LTR transposons (SanMiguel et al. 1998) (Sup. Fig. 4; Sup. Fig. 15). We see evidence of rare long-range duplications of NLRs, likely mediated by copy-paste TE activity: at least 12 neighbourhoods contain NLRs or NLR pseudogenes in only a few accessions, are TE-rich, and are not near any other NLR neighbourhoods (Sup. Fig. 16). While these could represent NLR neighbourhoods that have lost NLRs in all but a few accessions, the ages of linked TEs suggest that these NLR neighbourhoods are most parsimoniously the result of copy-paste activity of TEs that moved partial or intact NLR genes. The presence of multiple homologous TE sequences is furthermore likely to provide a substrate for NLR gene duplication that is independent of TE copy-paste activity, but rather the result of localised processes such as replication slippage and illegitimate recombination. There are many examples where TE seems to be directly responsible for the removal of NLRs including ADR1 an nominally essential helper NLR that is missing from one of the accessions (Sup. Fig. 20)

## Discussion

For three decades, studies of plant NLRs, a major class of immune receptors, have often stressed their evolutionary fluidity. This emphasis on structural and sequence variability of NLRs is not surprising, given that the functional identification of NLRs in many cases began with naturally occurring genetic variation. More recent work has suggested that extreme intraspecific diversification is a hallmark of a special class of hypervariable NLRs, while many other NLRs are evolutionarily much more stable. Here, by considering both structural and sequence variation, we show that highly conserved NLRs and hypervariable NLRs represent merely the extremes of a continuum, and that NLR diversity itself is much more complex than previously recognised.

### Context uncovers the extent of NLR evolution

Previous studies identified highly complex regions of the genome, a subset of which contain NLRs (Jiao and Schneeberger 2020), or studied NLRs either without genomic context (Van de Weyer et al. 2019), or in the context of a single individual (Sutherland et al. 2024). By exhaustively annotating intact and degraded NLRs in pangenomic neighbourhood context, we highlight that NLR loci vary considerably in structural complexity, with the majority of NLRs existing in relatively structurally conserved regions. A major surprise was that, when NLR neighbourhoods are structurally complex, the complexity is driven by the NLR genes themselves, rather than NLRs being passengers in regions of the genome that are generally complex.

The highly dynamic nature of NLRs in structurally complex neighbourhoods is reminiscent of the vertebrate adaptive immune system (Boehm and Swann 2014), with the proviso that NLR diversity plays out at the population level instead of within individuals. It is obvious that somatic recombination and hypermutation of the loci encoding vertebrate immune receptors is the outcome of selection for a maximally effective immune system of the individual (Giorgetti et al. 2023). It is difficult to think that the same logic does not apply to plant immune receptors, even though the generational scale of diversification is a different one. Indeed, several – though not all – of the structurally most complex NLR neighbourhoods harbour genes that are known to confer resistance to a highly co-evolving obligate biotrophic oomycete pathogen, *Hyaloperonospora arabidopsidis* (Holub 2001; Nemri et al. 2010). There is, however, a fine line between adaptation and maladaptation, and several of these neighbourhoods also harbour genetic loci that can cause autoimmunity (Chae et al. 2014; Jiao and Schneeberger 2020; Prigozhin and Krasileva 2021).

### The diversity of NLRs cannot be generalised

While a few NLRs are at the same time highly diverse across multiple axes, many more NLRs have elevated diversity according to only one of our seven metrics. Focusing on confirmed resistance genes, we find that they span the spectrum of diversity across all axes. Some resistance genes that are co-evolving with pathogen effectors that they directly recognise achieve extreme diversity in many axes, with many copies and a high degree of sequence variation (e.g. *RPP1* and *RPP4/5*), what is often seen as the prototypical pattern for resistance genes (van der Biezen et al. 2002; Yi and Richards 2007; Prigozhin and Krasileva 2021). Most resistance genes, however, have elevated diversity in only some axes: *RPP13* exhibits a large degree of variation in its primary sequence, but is otherwise a single-copy gene with comparatively low diversity along all other axes (Bittner-Eddy et al. 2000). The sequence of *RPP7* is relatively conserved across accessions, with limited copy number variation, but with many independent inactivating mutations across the wider population. While one can derive simple metrics for NLR diversity (Sutherland et al. 2024), the use of multiple metrics clearly provides a much more nuanced picture of NLR diversity. That said, single metrics have promise in highlighting candidates for bioengineering (Brabham et al. 2024). While it is important to understand the constraints of NLR diversification imposed by mutational mechanisms, a pathogen does not care if its receptor is disabled by point mutation, deletion, or TE insertion, only that it can now invade unrecognised.

### The many axes of NLR diversification

The diversity observed at any locus is the product of local mutation rates and the specific selective landscape acting on a locus, and genomic context improves our understanding of the processes generating and maintaining NLR diversity (Van de Weyer et al. 2019; Lee and Chae 2020; Prigozhin and Krasileva 2021; Sutherland et al. 2024). For example, gene duplications occur more readily where local segmental duplications already exist (Reams and Roth 2015). Such duplicated copies of an NLR gene could allow additional recognition specificity to evolve, or could pose increased costs, for example illegitimate oligomerization that impedes proper molecular functionality (Stirnweis et al. 2014). Whether these differences result from the underlying DNA sequence biassing mutational processes, or whether it is imposed through selection is less easy to determine, but we can surmise a likely answer in many cases.

While NLR loci are well-known hotspots of gene duplication, inactivation, and loss (Michelmore and Meyers 1998), relatively few NLR orthogroups are found in multiple neighbourhoods, indicating that most copy number expansion occurs locally (Baumgarten et al. 2003), although we observe at least 12 apparent TE-mediated distal duplications of NLRs to neighbourhoods. The causes of NLR inactivation by premature stop codons or frame shifts generally reflect the spectrum of mutations disabling non-NLR genes in *A. thaliana* (Monroe et al. 2018; Monroe et al. 2021). Taken at face value, this would seem to indicate that inactivating mutations or pseudogenization are more common in NLR neighbourhoods than elsewhere because of a greater tolerance for NLR mutations. The punctuated nature of pathogen selection may lead NLR neighbourhoods to frequently experience episodes during which they are less constrained by purifying selection, and consequently we see mutations that would otherwise have been purged, especially when intact NLRs are mildly deleterious in the absence of pathogens.

While NLR neighbourhoods are enriched for TEs, TEs play only a minor direct role in both gene duplication and pseudogenisation events, even though they tend to be young and turn over quickly in NLR neighbourhoods. We do not know whether this is due to increased rates of TE insertion, or to increased persistence of TEs, possibly due to fluctuating selection dynamics whereby TE insertions are tolerated in the absence of a pathogen, then swept to high frequency as passengers on selectively advantageous NLR haplotypes in the presence of a pathogen. It is also possible that TEs are positively selected in some NLR neighbourhoods as they impact the regulatory landscape (Zervudacki et al. 2018; Panda and Slotkin 2020). TE insertions likely play some role in immune system maintenance: a relatively simple example is the *CHS3/CSA1* paired NLR locus, where two distinct haplotypes are maintained across the population (Yang et al. 2022), likely aided by the presence of highly divergent intergenic TEs that prevent recombination between the two NLRs. We also frequently observe multiple copies of NLRs and adjacent TEs within an NLR neighbourhood, possibly due to segmental duplication by replication slippage or illegitimate recombination, possibly accelerated by additional TE copies (Reams and Roth 2015).

Alternative splicing increases transcript diversity, and there are examples of isoform variation generating immune diversity in plants (Yang et al. 2014; Kufel et al. 2022). We found that individual NLR genes vary in isoform diversity, but that NLR genes from the same orthogroup have similar isoform diversity. While our study design does not reveal changes in isoform diversity during the course of infection, the observation of frequent instances of alternative splicing make it likely an additional important axis of NLR variation. NLR expression has been shown to vary among accessions (Gan et al. 2011) and across environmental clines (MacQueen and Bergelson 2016), further highlighting the need to consider regulation of expression in studies of functional NLR diversification. Similarly, structurally-complex loci around NLRs are highly diverse in methylation patterns across accessions (Kawakatsu et al. 2016), highlighting the need to consider both inter-accession variation, as well as the dynamics of methylation around NLRs over the course of pathogen invasion and defence, which our single time point data cannot evaluate.

To efficiently explore present-day diversity, we deliberately chose accessions as single representatives of regional populations, between which recent gene flow and therefore recombination is limited. An obvious next step will be a careful comparison of NLR neighbourhoods across multiple populations with closely related individuals. This will allow, for example, assaying population recombination rates, and to determine exactly how they are affected by structural variation (Choi et al. 2016).

### Plants have adaptive immunity not as individuals, but as populations

We posit that “diversity in diversity” allows for evolutionary innovation at multiple evolutionary speeds, enabling plants to keep pace with pathogens, in which rapid genetic change is fueled by processes that range from two-speed genomes to formation of minichromosomes and horizontal gene transfer (Möller and Stukenbrock 2017; Bertazzoni et al. 2018). Plants do not take a single approach to generating diverse immune receptor complements, and any investigation of their evolution must consider all relevant axes along which diversification can occur. While individual plants do not have adaptive immune systems in the vertebrate sense, like vertebrates, plants must survive the onslaught of a highly diverse set of fast evolving pathogens. To persist, annual plant populations must constantly balance the detection of novel pathogen signals with fitness costs that come from a hyper-active immune system. Critically, as long as enough of the plants in a population have the right immune genes to survive that year’s wave of pathogens with only a minority of individuals having either too little or too much immunity, the population persists. An interesting question for the future is how all this plays out in long-lived plants such as trees, which will encounter over their lifetime a much greater diversity of pathogens than an ephemeral annual herb such as *A. thaliana*.

## Methods

### Accession selection and plant growth

We selected 17 accessions of *Arabidopsis thaliana* based on geographic stratification, seed availability, and previously identified haplotype sharing groups (Shirsekar et al. 2021). We grew plants in a potting mix at 23°C with 16 hours light and 65% humidity until 30 days old. We harvested whole rosettes after 30 hours of dark treatment to reduce starch, and stored them at -80°C.

### HMW DNA Extraction

To extract high molecular weight (HMW) nuclear DNA with minimal organellar contamination, we first isolated nuclei. Approximately 40 grams of plant tissue was ground to an ultra-fine powder with a mortar and pestle cooled with liquid nitrogen. Working at 4°C, we resuspended about 20 grams of ground tissue in 200mL of nuclei isolation buffer (NIB), and gently stirred for 15 minutes. The solution was filtered through Miracloth, and incubated with 10 mL NIB-Triton20 for 15 minutes. We centrifuged the solution to collect the nuclei and washed the pellet twice with NIB-Triton20, and finally centrifuged again to collect the final clean nuclei pellet.

We extracted HMW DNA by lysing nuclei in G2 lysis buffer at 37°C, treating with RNAse for 30 minutes at 37°C and then proteinase K at 50°C for 3 hours. We centrifuged this lysate at 4°C to remove debris. We eluted and precipitated the HMW DNA in the genomic tip device, then physically separated clumped DNA with a sterile glass hook and eluted in Qiagen EB buffer. We left the eluted HMW DNA at room temperature until it fully dissolved and stored it at 4°C. We measured HMW DNA fragment sizes with the Femto® Pulse system (Agilent), and quantitated DNA with the dsDNA-HS Qubit Fluorometer kit (Life Technologies). Typical yields were 30-80 µg of HMW DNA, with median fragment lengths of over 60-80 kbp.

### PacBio HiFi Library Preparation

We prepared the extracted genomic DNA for PacBio Circular Consensus Sequencing (CCS) with the “Procedure & Checklist - Preparing HiFi SMRTbell Libraries using SMRTbell Express Template Prep Kit 2.0 (PN 101-853-100 Version 01 (September 2019))” protocol, with the following modifications. We sheared the genomic DNA with the Megaruptor 2 device (Diagenode) to a target size of 20 kb to 25 kb. To reduce clogging of the Megaruptor hydropore, we sheared 15 µg HMW genomic DNA in a volume of 700 µl to an average size of between 15-20 kb. We purified sheared DNA with AMPure PB Beads (Pacific Biosciences), and checked the concentration and size with the Qubit Fluorometer and the Femto Pulse System. We used the Express Template Prep Kit 2.0 for library construction, except we used double the suggested input DNA (10 µg of sheared DNA) and accordingly increased all reagent volumes two-fold. The library was size selected to >15 kb using a BluePippin instrument. Immediately before sequencing on a Pacific Biosciences Sequel II instrument, we performed a final bead cleanup, annealed the templates with Sequencing Primer v2 and bound it to the sequencing polymerase.

### HiFi Read generation and DNA Methylation estimation

We generated HiFi consensus reads from raw read files using the PacBio Circular Consensus Sequencing tool (v6.0.0) and bam2fasta, keeping reads ≥ 10 kb and with average quality ≥Q20 for genome assembly. We re-generated HiFi consensus reads with kinetics information with ccsmeth call_hifi (Ni et al. 2023). We generated an Illumina bisulfite sequencing (BS-seq) library as described (Mencia et al. 2023) for a single accession (at9852) using a portion of input DNA from which PacBio CCS reads had been generated. We used Bismarck (Krueger and Andrews 2011) to generate estimates of cytosine methylation from the BS-seq reads. We trained a ccsmeth model (v0.3.0; (Ni et al. 2023)) to call methylation based on HiFi read kinetics, using the Bismarck methylation estimates as ground truth. Briefly, genomic positions with 100% methylation and at least 6x BS-seq read depth were considered methylated and those with 0% methylation were considered non-methylated. During cross-validation, our fine-tuned ccsmeth CG model had 98% accuracy, but CHG and CHH sites could not be accurately predicted. We then used ccsmeth to estimate methylation status for each HiFi reads for all accessions. We used methylartist (Cheetham et al. 2022) to aggregate read-level methylation data for individual cytosines in each genome.

### Genome assembly and quality assessment

We assembled reads into primary contigs with hifiasm (v0.15.4-r343) (Cheng et al. 2021). We mapped HiFi reads back to primary contigs with minimap2 (Li 2018; Li 2021) and samtools (Li et al. 2009; Danecek et al. 2021), and searched each contig against the NCBI’s non-redundant protein database with diamond blastx (Buchfink et al. 2021). We used blobtools (Laetsch and Blaxter 2017) to combine these data and generate summary plots that we used to identify contaminant sequences, which we removed with seqtk (Li 2008). To increase the accuracy of scaffolding, we used an optical map (produced by Bionano Genomics) of accession at9852. We scaffolded contaminant-filtered primary contigs of at least 100 kb by aligning them to the Bionano map of at9852 using ragtag scaffold (v2.0.1; (Alonge et al. 2022)). We assessed the completeness and correctness of each de novo assembly with BUSCO (v4.0.6) (Simão et al. 2015) using the odb10_embryophyta database (Zdobnov et al. 2021). We calculated continuity, GC content, overall assembly length and reference coverage of assembled contigs with Quast (v5.0.2) (Gurevich et al. 2013). We detected structural variation between each scaffolded genome and the *A. thaliana* reference genome TAIR10 (The Arabidopsis Genome Initiative 2000) with SyRi (v1.3) (Goel et al. 2019). The assembly of at6035 was flagged as a likely mix of at least two accessions and therefore excluded from subsequent analyses.

### *Ab initio* and liftoff gene annotation

We created initial gene annotations using both homology evidence and statistical prediction. We created *ab initio* gene predictions with Augustus (Nachtweide and Stanke 2019), using a BUSCO-refined *A. thaliana* model similar to the BREAKER workflow (Brůna et al. 2021). We used liftoff (Shumate and Salzberg 2021) to transfer the Araport11 (Cheng et al. 2017) annotation of the Col-0 reference accession to homologous regions of each assembly. We also produced a protein domain annotation, for all putative gene models using InterProScan (interproscan-5.51-85.0 (Jones et al. 2014)). We then translated these matches into GFF files using pfam2gff.py (https://github.com/wrf/genomeGTFtools/blob/master/pfam2gff.py).

### Gene annotation with transcript evidence

To produce transcript evidence for gene annotation, we infected 10-day old seedlings of each of the 17 accessions with three isolates of the specialist oomycete pathogen *Hyaloperonospora arabidopsidis* (166-4, 495-1, 527-3). We harvested two replicates each at 2 hours, 24 hours, and 4 days post infection, and extracted RNA with a silica column-based protocol (Yaffe et al. 2012; Yuan et al. 2023). We quantified the RNA using a Nanodrop instrument and pooled all samples for each accession in an approximately equimolar fashion. We prepared Iso-Seq libraries using the “Preparing Iso-Seq libraries using SMRTbell® prep kit 3.0” protocol. Libraries were individually indexed using SMRTbell barcoded adaptors during the adapter ligation step of the library prep protocol. The libraries were multiplexed and sequenced across 3 SMRT cells (8-10 libraries per SMRT cell) using the Sequel II binding kit 3.1 recommended for libraries with fragments shorter than 3 kb. The ten libraries with the lowest yields were re-pooled and re-sequenced on a fourth SMRT cell.

To predict isoforms from the Iso-Seq reads we used a custom pipeline that can handle multiple reference genomes (https://github.com/Lnve/HYDRA). Briefly, Pacbio CCS reads were demultiplexed and barcodes removed using Lima (2.6.0) with the Iso-Seq mode. We clipped poly-A tails using refine (3.8.1) and removed apparent concatemers. We merged samples that were sequenced twice with samtools (1.16.1). We mapped reads to the respective reference genome using minimap2 (2.17-r941 -ax splice:hq; (Li 2018; Li 2021)) and collapsed the reads into predicted isoforms using TAMA (tc_version_date_2020_12_14; (Kuo et al. 2020)). We set the 5’ threshold (-a 10) to 10 bp, the 3’ threshold to 5 bp (-z 5) and did not allow for any differences at the splice junctions (-m 0), resulting in variation at the ends, but capturing all possible splice junctions of each gene. Only sequences with 99% identity were collapsed (-i 99). We then updated the initial gene predictions from AUGUSTUS with PASA (2.5.2; (Haas et al. 2003)) using default settings. We predicted open reading frames for all isoforms using transdecoder (5.7.0; (Haas et al. 2017)) with the complete ORF setting.

### Evidence-weighted gene prediction

To integrate *ab initio*, liftoff and RNA supported gene predictions, we combined all predictions into non-overlapping loci likely representing the same underlying gene(s). We used NLRannotator (Steuernagel et al. 2020) and NLRtracker (Kourelis et al. 2021) to find NLR remnants or NLR genes missing from our annotated genes. We devised a decision tree (see Sup. Fig. 3) to categorise genes according to the amount and source of consistent evidence. Where annotators agreed on the coding sequence and exon structure of a gene, we selected the most evidence-rich annotation (PASA > liftoff > augustus). Where RNA-based annotation disagreed with *ab initio* and/or liftoff prediction, or where RNA evidence was missing, we manually curated NLR annotations.

The manual curation process involved examining the locus in both webapollo (Lee et al. 2013) and igv-js (Robinson et al. 2023), and assessing which evidence track contained the valid gene model with the most canonical NLR domain structure supported by both RNA and homology evidence. Manual annotation was mostly needed for a small set of predictions where the Iso-seq evidence overlapped with multiple *ab initio* and liftoff annotations, or where there were multiple liftoff annotations for a single Augustus annotation and vice versa. Where we encountered valid open reading frames encoding NLRs with non-canonical domain structure, we used Iso-Seq evidence as the putative gene model. In the absence of Iso-Seq evidence, if putative annotations supported differing numbers of genes, we examined the domain structure of these ORFs to determine whether the “split” or “conjoined” putative annotation was likely more accurate, informed by the domain structure of homologous sequences from other accessions. For example, if liftoff of Araport11 predicted two gene models but the Isoseq and the domain structure predicted a single NLR with a TE insertion, we selected the gene model predicted using the Isoseq. We used a custom script to parse the manual annotation decisions and output a composite GFF. GFF entries were adjusted for some truncated pseudogenes, where the extent of the mRNA boundaries were lengthened to encompass the full 3’ UTR of the truncated gene. Finally, we combined the outputs for each accession, which we then sanitised for a variety of common minor problems using both AGAT (Dainat et al. 2023) and custom tools (Murray 2024a). Where multiple transcripts were predicted for a gene model, the transcript with the 5’-most start codon and then longest CDS was chosen as representative. Genes were labelled as pseudogenes if there was a liftoff annotation that did not have a valid ORF. Proto-pseudogenes/truncated genes were labelled as part of the manual NLR annotation process, usually where only one mutation interrupted the NLR ORF, often with Iso-seq evidence for a transcript with a long 3’ UTR downstream of the mutation truncating the ORF.

### Isoform diversity

We summarised isoform variation based on the ORF structure of each isoform (i.e. collapsing isoforms with the same open reading frame) using SQUANTI3 (Pardo-Palacios et al. 2024). We calculated the isoform diversity for each gene as Simpson’s index of diversity (1-Simpson’s D; 1=maximal diversity). As observed isoform diversity is limited by the number of reads sequenced, we also recalculated Simpson’s index of diversity only for genes that had at least 10 Iso-seq reads. As we did not explicitly enrich for intact 5’ caps, we also re-calculated isoform diversity disregarding isoform variation at the 5’ end to account for potential transcript degradation.

### Repeat annotation

We annotated satellite repeats such as telomeres, centromeres and rDNA clusters by homology to a library of consensus sequences from the reference accession Col-0 (Rabanal et al. 2022) using RepeatMasker (Chen 2004) (v 4.0.5) with the following parameters: `-e ncbi -s -a -inv -xsmall -div 40 -no_is -nolow -cutoff 200 -norna`. To annotate transposable elements (TEs) in each genome, we first used EDTA_raw.pl (v.1.9.7) (Ou et al. 2019) to annotate LTRs with LTRharvest (v1.5.10) (Ellinghaus et al. 2008), LTR_FINDER_parallel (v1.0) (Ou and Jiang 2019) and LTR_retriever(Ou and Jiang 2018), helitrons with HelitronScanner (v1.0)(Xiong et al. 2014), and TIR elements with TIR-Learner (v1.23) (Su et al. 2019) and MITE-Hunter (v1.0)(Han and Wessler 2010). We then merged all 90 chromosomes from the 18 accessions, combined the raw output and proceeded with the rest of the EDTA pipeline, adding the current TE library from TAIR10. Originally 18 genomes were included in this dataset but we could not generate IsoSeq expression evidence for at6137 so it was not included in any analysis of NLR diversity and evolution. The end result was a combined TE library and GFF annotation of TEs common for all the accessions.

To refine these automated annotations and to detect novel repeat families, we used additional steps: To mitigate rampant misannotation of tandem repeats as TEs, we intersected EDTA annotation with the independent satellite annotation of centromeres, telomeres and rDNA and removed EDTA-annotated TEs overlapping >20% with satellite repeats. We removed repeats not assigned to a known repeat family, as they were predominantly either artefacts of the joint analysis of all 18 genomes, or unidentified satellite repeats.

To increase the confidence of the *de novo* helitron annotations, we considered whether a TE family had at least one intact member, and whether there was a Rep/Hel protein domain in at least one member (using RexDB Viridiplantae v3.0 (Neumann et al. 2019). Finally, we ran EAhelitron (Hu et al. 2019) to reannotate helitrons and report whether a given TE copy intersected with both EDTA and EAhelitron.

The EDTA pipeline did not assign a TE family to several TE instances because they were single copy, likely because we had run the raw EDTA module independently for each genome. To correct this, we associated elements with known TE families via BLAST matches following the 80/80/80 rule. We clustered the remaining TE instances that did not correspond to a known TE family following the 80/80/80 rule using CD-hit (Li and Godzik 2006).

Clusters with at least two copies were assigned new TE family names and their corresponding TE models were incorporated into the TE library of the pangenome. Finally, we used TEsorter (Zhang et al. 2022) to assign all LTR families in our TE library to known clades using RexDB Viridiplantae v3.0 (Neumann et al. 2019). Finally, we estimated the age of intact LTRs by aligning their two LTR ends and calculating the pairwise distance between the two ends using the dist.dna function of the R package “ape” (Paradis and Schliep 2019) with the model “K80” and translating it to millions of years using the mutation rate calculated in (Ossowski et al. 2010).

### TE gene annotation

To distinguish *bona fide* TE genes from non-TE genes, we devised the following decision tree (Sup. Fig. 21). We combined all TE and non-TE genes produced by the annotation (together referred to as putative genes from now on), and used Diamond v2 (Buchfink et al. 2021) to compare their protein products against a curated repeat database of TE-related protein domains (RexDB Viridiplantae 3.0 (Neumann et al. 2019)). We overlapped all putative genes with a collapsed version of the TE annotation without helitrons, merging all annotated TE copies, minus helitrons, that overlap or are within 100 bp of each other. We classified putative genes as TE genes if they overlapped at least 20% with merged TEs and had a RexDB hit. In addition, we classified putative genes as TE genes if they did not have a hit against RexDB but overlapped at least 90% with merged TEs. We compared all putative genes with a RexDB hit with a merged helitron annotation, and classified those with more than 20% overlap as TE genes. We classified as “TE-like protein genes” all other putative genes with a RexDB hit but without overlap with the TE or heliotron annotations. All remaining genes without a RexDB hit and without overlap with merged TEs or helitrons as protein coding genes.

### Orthogroups, synteny, and phylogenetic inference

To identify NLR sequences that represent the same gene, we first extracted the protein sequences of all genes with a valid ORF (excluding pseudogenes but including truncated genes) using AGAT (Dainat et al. 2023). We also extracted the nucleotide sequences for all genes (including pseudogenes and pseudogenic regions). We clustered all protein sequences across the 17 accessions using OrthoFinder2 (Emms and Kelly 2019) as an initial first pass with default settings. This resulted in 35,697 orthogroups (OGs), of which 249 contained NLR proteins. We confirmed the global synteny of the 17 genomes using Genespace (Lovell et al. 2022) (Sup. Fig. 2).

Many diversity metrics are heavily affected by how orthogrouping is performed, which can be quite arbitrary, particularly for NLRs with both recent and not so recent copy number variation. We therefore defined broad orthogroups (OGs) along with finer orthogroups (OG70s). For OGs, we first conducted an all-by-all protein blast (BLASTP (Camacho et al. 2009)) to ensure that sequences had not been mis-assigned by OrthoFinder and to determine whether any OGs needed to be merged. If only a few sequences in an OG had close hits (≥ 85% amino acid sequence identity) to another OG, we moved these sequences to the other OG. If most sequences of an OG had a close hit with sequences from another OG, we merged the OGs. This procedure reduced the number of OGs from 249 to 204, and led to OG reassignment of 24 sequences. To produce final OGs representing sequences of the same NLR gene, we clustered sequences within OGs using CD-hit (Li and Godzik 2006) with a 70% sequence similarity threshold to produce OG70s. By definition, all members of an OG70 belong to the same OG. We aligned sequences in each OG70s using mafft –auto (Katoh et al. 2005) calculated average pairwise distance using custom script - p-distance_script_v3.py and Shannon entropy using Blindschleiche (blsl entropy (Murray 2024a)).

We assigned pseudogenes to OGs using blastx (BLAST+ (Camacho et al. 2009)), aligning all NLR proteins to the nucleotide sequences of the pseudogene regions. A pseudogene was assigned to the orthogroup of the protein that best aligned to each pseudogenic region. We did not search against the entire protein complement as the dataset contains TE proteins, many of which exist inside the pseudogenes.

We constructed gene trees for orthogroups (OGs and OG70s) of interest (e.g. the orthogroup containing ADR2, Fig5 b). We aligned nucleotide sequences for the entire genes using MAFFT (Katoh et al. 2005) and reconstructed a Maximum likelihood phylogeny using IQtree (Minh et al. 2020) and visualised the tree using ggtree (Yu 2020).

### Pangenome graphs

We induced a graph of each of the five chromosomes from all assemblies with PGGB (Garrison et al. 2023) using the parameters recommended for *A. thaliana* (-s 5000, -p 95, -n 18, -k 47). We visualised graphs and subgraphs with odgi viz (Guarracino et al. 2022). We calculated local graph complexity with a graph-walking metric: for each node in the graph counted the total number of unique nodes that could be visited with a specific number of steps without back-tracking and ignoring self-loops. We walked each assembly’s path through the graph, transferring node-level statistics to assembly-specific genome coordinates. The functionalities are implemented in the tool ’raugraf’ (Murray 2024b).

### Definition of NLR-dense neighbourhoods

We defined NLR-dense gene neighbourhoods (“NLR neighbourhoods”) with a semi-automated pangenome graph traversal algorithm. For each chromosome, we find the position of an NLR in the pangenome graph. We then traverse outward until we find at least five nodes with at least 100 bp of syntenic, single-copy sequence shared across all accessions. (Fig. 1b). These syntenic anchors from the left and right borders of a region with at least one NLR in at least one accession. We then define a NLR neighbourhood in each accession by finding the corresponding genome coordinates of these syntenic anchor nodes with odgi. We repeat this for all NLRs in all accessions, skipping those already covered by a neighbourhood, until all annotated NLRs are contained in neighbourhoods.

We removed two neighbourhoods that contained only a single incomplete NLR fragment across the pangenome, in both cases a partial TIR sequence appears to be inserted into pericentromeric TE repeat clusters.

### Genotyping with data from the 1001 Genomes project

To determine the species-wide frequency of mutations that disrupt the coding sequence of NLR genes, we applied the Acanthophis variant calling pipeline (Murray et al. 2024) to short reads from the 1001 Genomes project (Alonso-Blanco et al. 2016). We removed low quality and adapter sequences from reads with AdapaterRemoval (Schubert et al. 2016) and mapped reads to the at9852 assembly with BWA MEM (Li 2013), calling variants with freebayes (Garrison and Marth 2012). We predicted variant consequences with bcftools csq (Danecek and McCarthy 2017; Danecek et al. 2021), using our final gene annotation. We calculated the total frequency of severe mutations (frameshifts, starts/stops gained or lost) as the sum of the allele frequency of all variants with a minor allele frequency above 1%, a variant quality above 1,000, and a total coverage within 5,000-50,000 (across 1,135 samples, equivalent to an average depth of approximately 5-50x).

### Data and Software Availability

While data and software are being publicly released, we have created an online portal to collate these releases available at https://dl20.coevolutionlab.org/datarelease/.

## Acknowledgements

We thank Gal Ofir, Markéta Vlkova-Žlebková, Miriam Lucke, Sebastian Vorbrugg, Yueqi Tao, Fabrice Roux, and Haim Ashkenazy for scientific discussions and/or commenting on the manuscript. We thank Katrin Fritschi and Anette Habring for assistance with preparing IsoSeq libraries. We thank Anton Schölkopf and the students of the 2021 PBC practical at the University of Tubingen for assistance with manual NLR annotation. Computations were performed at the Max Planck Computing and Data Facility, Garching, Germany. We thank Marie Skłodowska-Curie Actions (to KDM), and DFG-TRR356 *PlantMicrobe*, the DFG-funded Excellence Cluster *Control of Microorganisms to Fight Infections* (CMFI), the Novozymes Prize of the Novo Nordisk Foundation, and the Max Planck Society (to DW) for research funding. LCT and KDM are co-first authors of this manuscript, and GS and DW are co-corresponding authors of this manuscript.

---

**Supplementary Figure 1:**
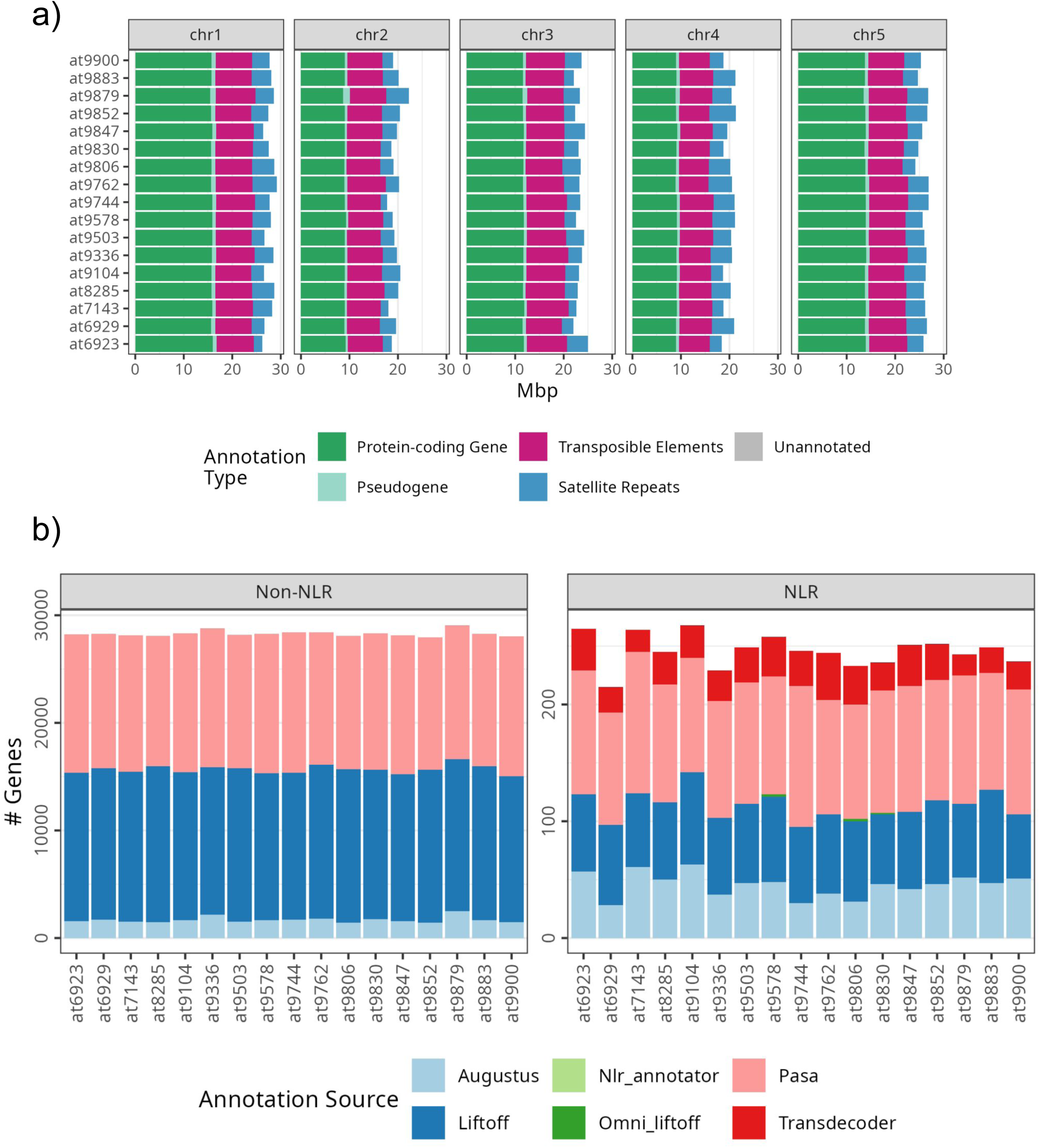
Annotation statistics. a) Breakdown of annotated sequence for each chromosome in each genome assembly. Most variation is in the satellite repeats. b) The annotation source for each gene for both the NLR and non-NLR genes. Only NLR genes were manually curated, and so only NLRs have gene entries from NLR annotator or Omni-liftoff.

**Supplementary Figure 2:**
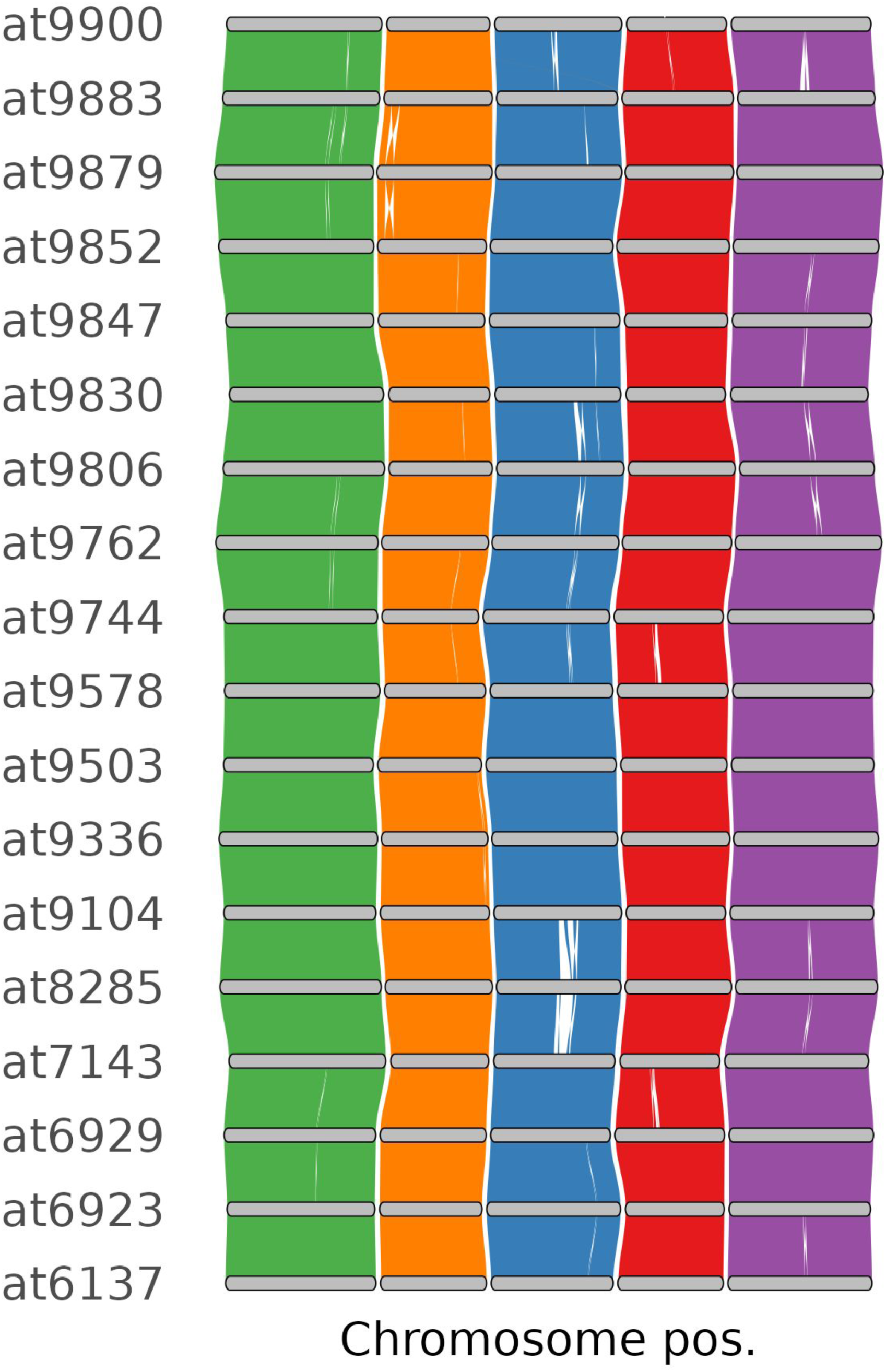
Global synteny is highly conserved across all 18 genomes, as estimated by Genespace which compares the relative position of each orthofinder orthogroup across all accessions.

**Supplementary Figure 3:**
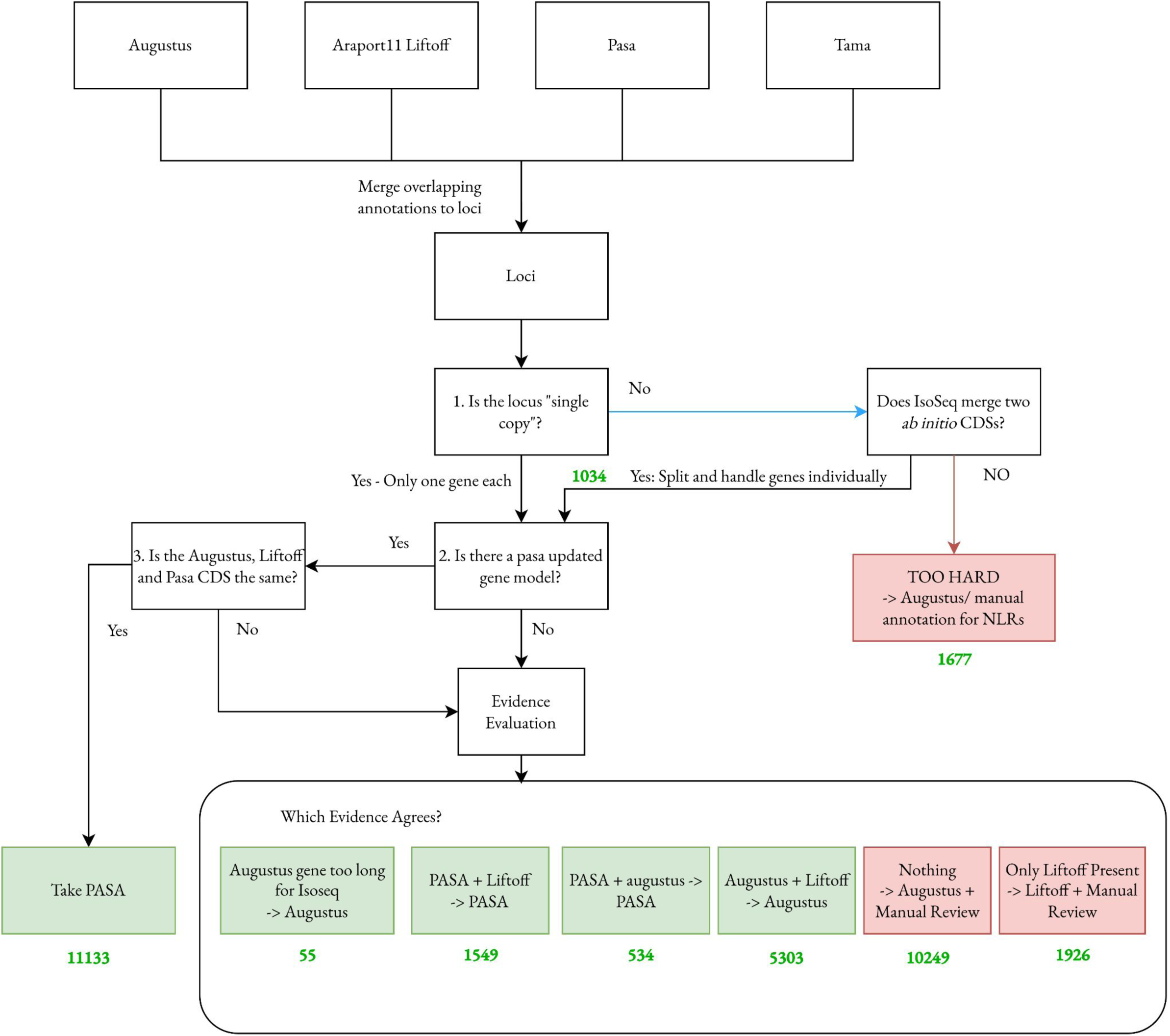
Gene annotation pipeline. We created candidate annotation tracks from ab initio (Augustus), homology (Liftoff), and expression evidence (Tama, Pasa). These candidate annotations were merged into overlapping loci. For loci with a single gene from all candidate annotators, we applied a decision tree as described below: briefly, we picked the most evidence rich valid gene model, bearing in mind caveats of each candidate annotation. For multi-gene loci, where IsoSeq evidence merged what were multiple independent coding sequences in all other annotations, we determined this was likely RNA read-through, and split genes into single CDS, which then entered the single-gene decision tree. For case where homology or ab-initio evidence disagreed as to how many genes were present, we took Liftoff, or in the case of NLRs, we manually decided which annotation to use (too hard). This automated pipeline was applied to all genes, but for NLRs, all genes were manually reviewed. For automated assignment, red boxes below represent “low confidence” annotations, which are flagged in the output GFF for non-NLRs, and should be taken as hypotheses at best. Green numbers represent the number of decision loci (not genes) that the respective automated decision was made for at9852

**Supplementary Figure 4:**
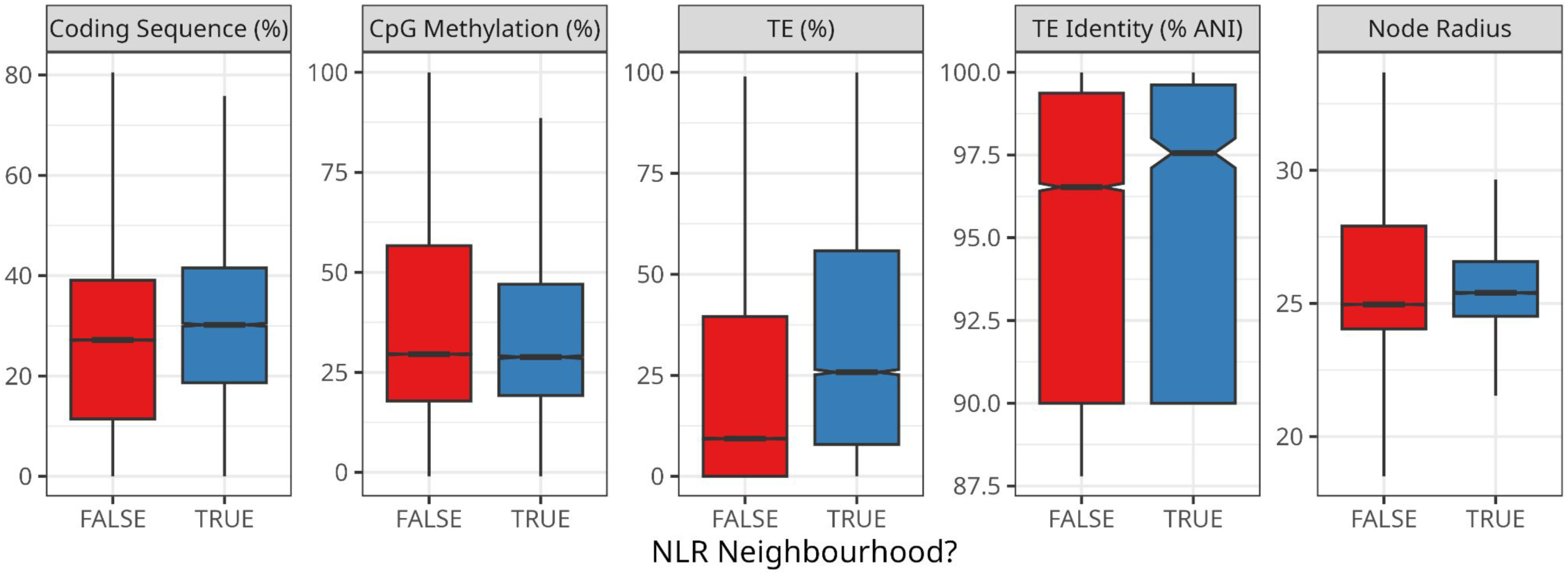
NLR neighbourhoods are broadly comparable to other genic parts of the A. thaliana genome, though are more TE dense. Genome windows of 10kbp that overlap with NLR neighbourhoods have similar steady-state CpG methylation as windows that do not overlap with NLR neighbourhoods. Windows that overlap NLR neighbourhoods contain significantly more annotated TEs, and their TEs are somewhat younger, as judged by each TE’s mean identity to the TE family centroid (TE Identity), and local pangenomic complexity (Node Radius) and the proportion of each window that is protein coding are marginally higher across NLR neighbourhoods compared to the background genome.

**Supplementary Figure 5:**
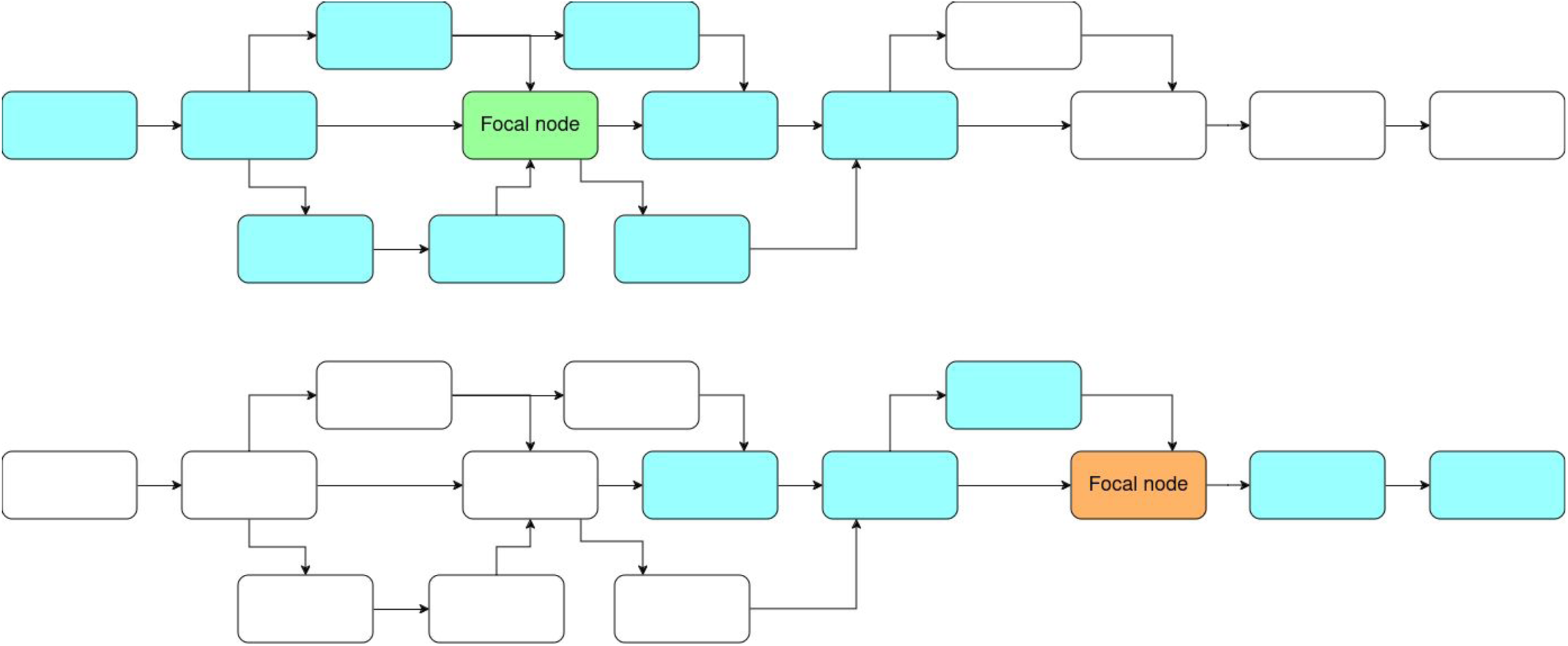
Simplified representation of the pangenomic complexity metric “Node radius”. For each node in the pangenome graph, we calculate how many distinct nodes are reachable within n steps from the focal node, regardless of the directionality of edges. The green focal node is located in a more complex region of the pangenome graph, and with n=2 steps in any direction we can visit 9 distinct nodes (blue). The orange focal node is located in a less complex region of the pangenomic graph, so with two steps we only visit 5 distinct nodes (blue).

**Supplementary Figure 6:**
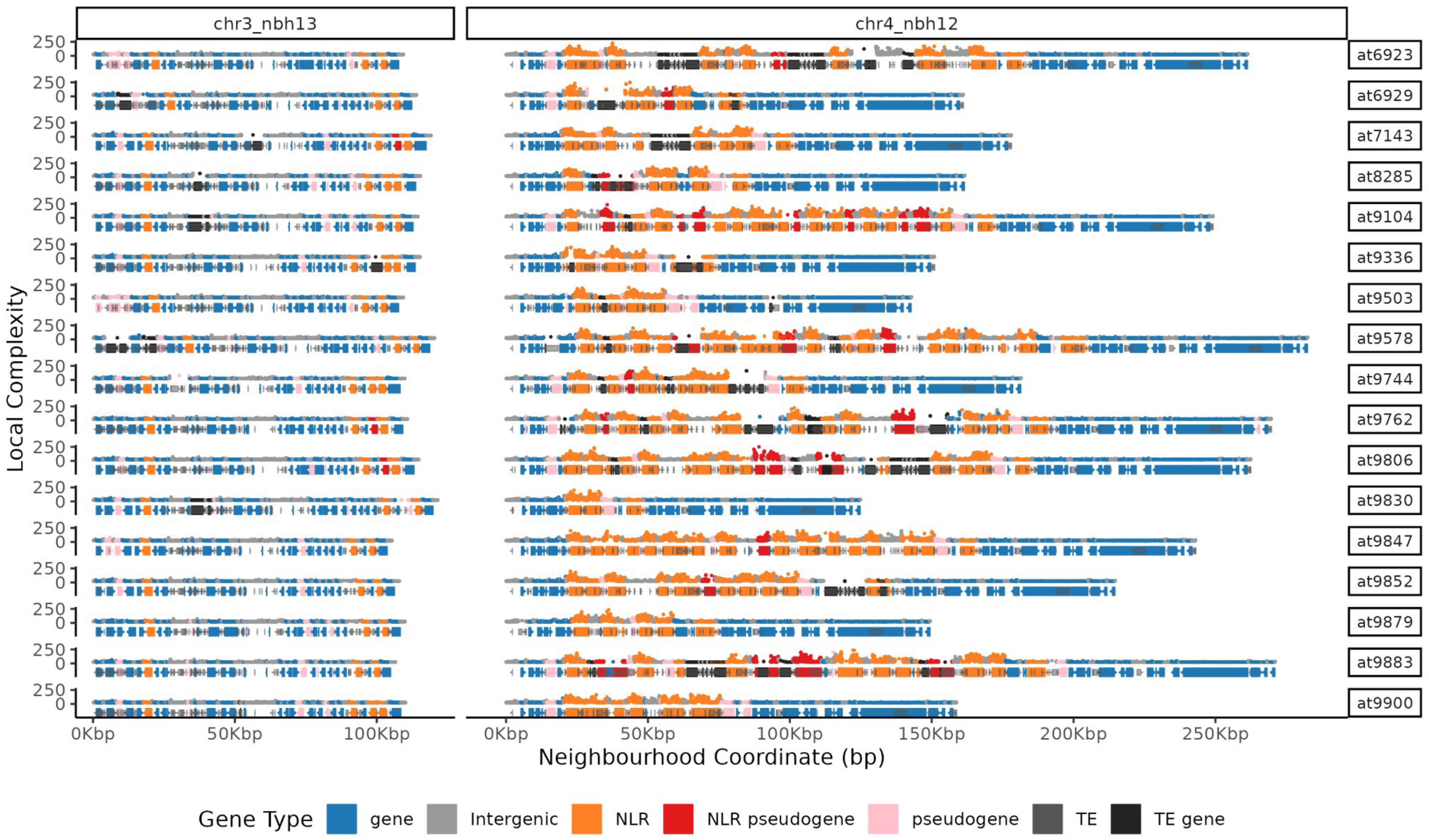
Local pangenomic complexity and gene annotations at two representative neighbourhoods, chr3_nbh13 contains *RPP13* and chr4_nbh12 contains *RPP4/5*. Local complexity is not particularly elevated in chr3_nbh13, which reflects the generally-syntenic nature of this region. In contrast, the extreme structural diversity of the *RPP4/5* locus (chr4_nbh12) greatly elevates our measure of local complexity, though specifically on NLR genes themselves.

**Supplementary Figure 7:**
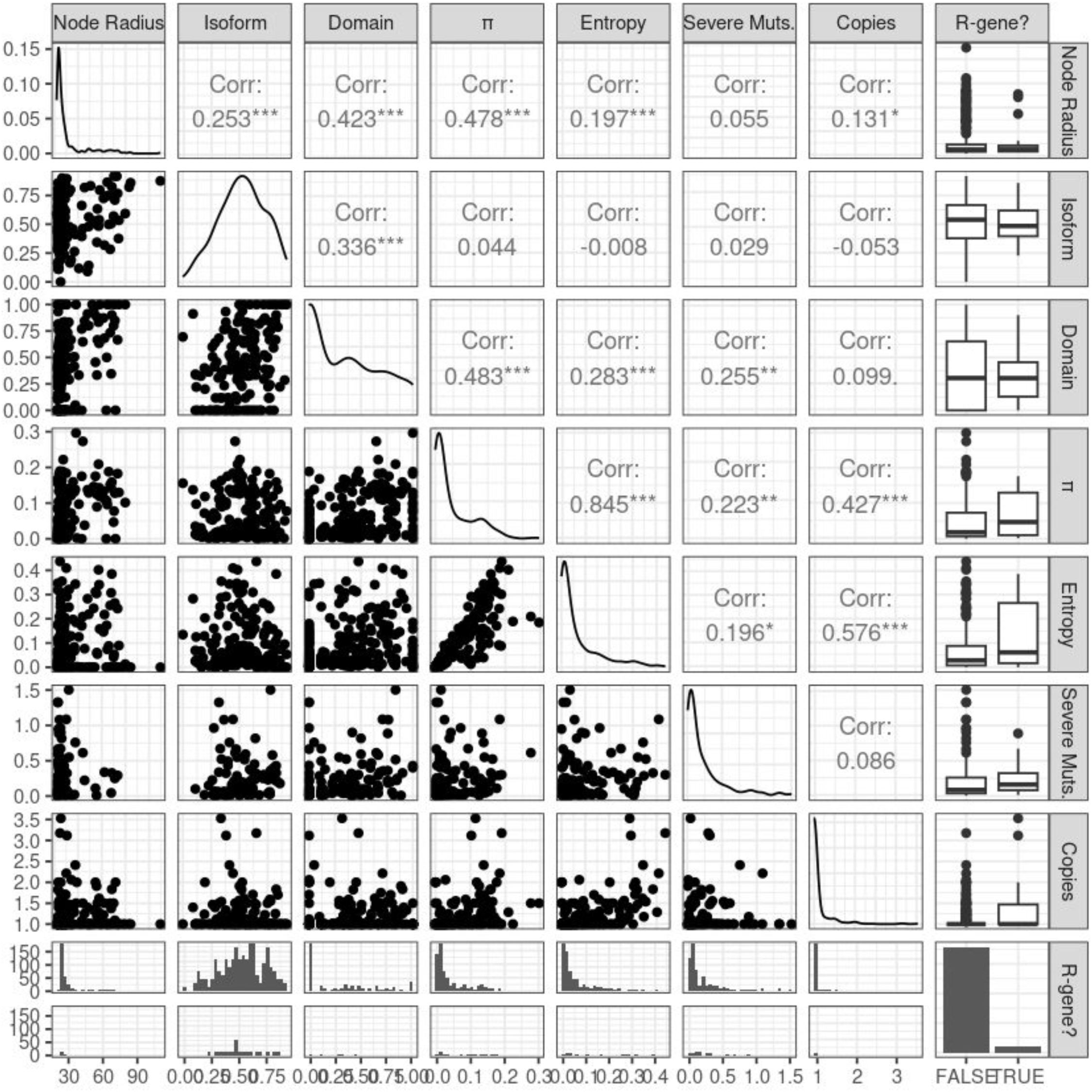
Correlation between metrics of NLR diversity at the OG70 level. Nearly all metrics are weakly correlated at best (0 < r < 0.5, i.e. R^2^ less than 25%), with only pi and mean shannon entropy showing a strong correlation (r=0.845; R^2^ = 71%).

**Supplementary Figure 8:**
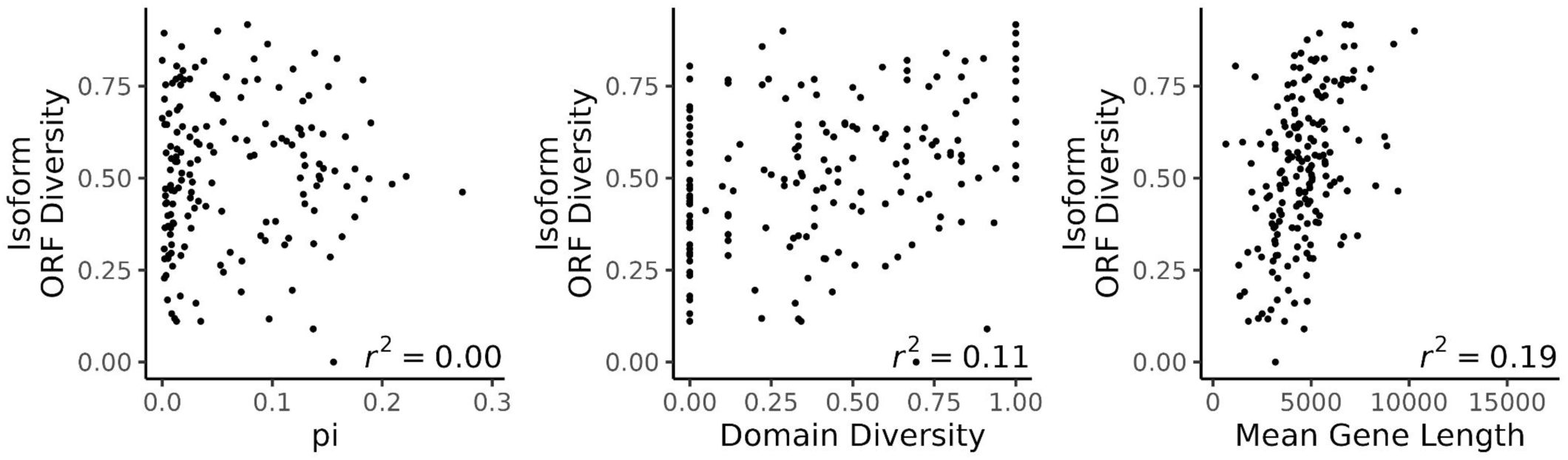
The relationship between Isoform diversity and several other NLR diversity metrics. Isoform diversity is not correlated with pi, but it is weakly correlated with domain diversity and mean gene length.

**Supplementary Figure 9:**
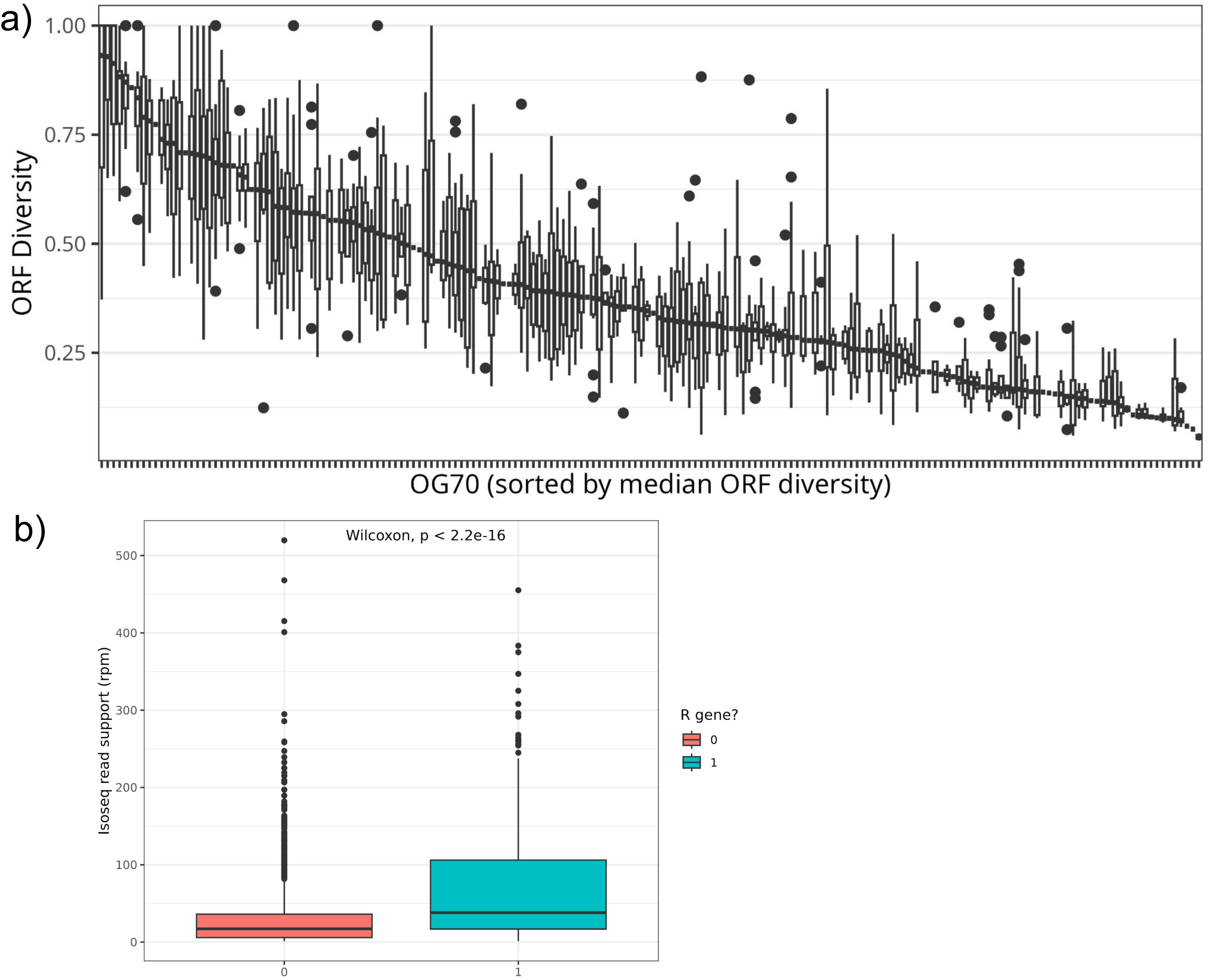
Isoform and Isoseq read coverage statistics. a) Isoform diversity varies across NLR orthogroups, but NLRs from the same orthogroup tend to have similar isoform diversities, b) know R gene NLRs on average have higher expression than other nlr genes in this dataset but there is lots of overlap.

**Supplementary Figure 10:**
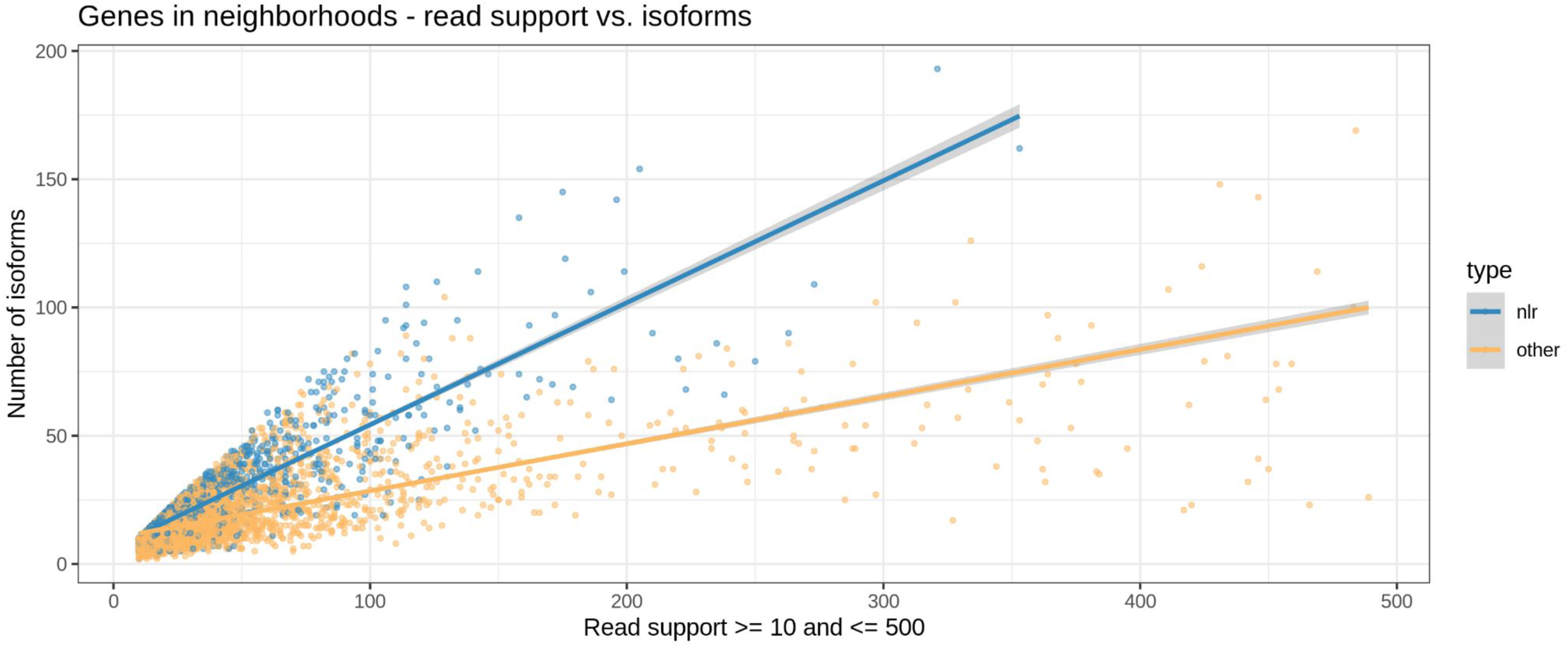
Of all the genes in NLR gene neighbourhoods, NLRs have more isoforms than non-NLR genes (only considering genes with at least 10 reads).

**Supplementary Figure 11:**
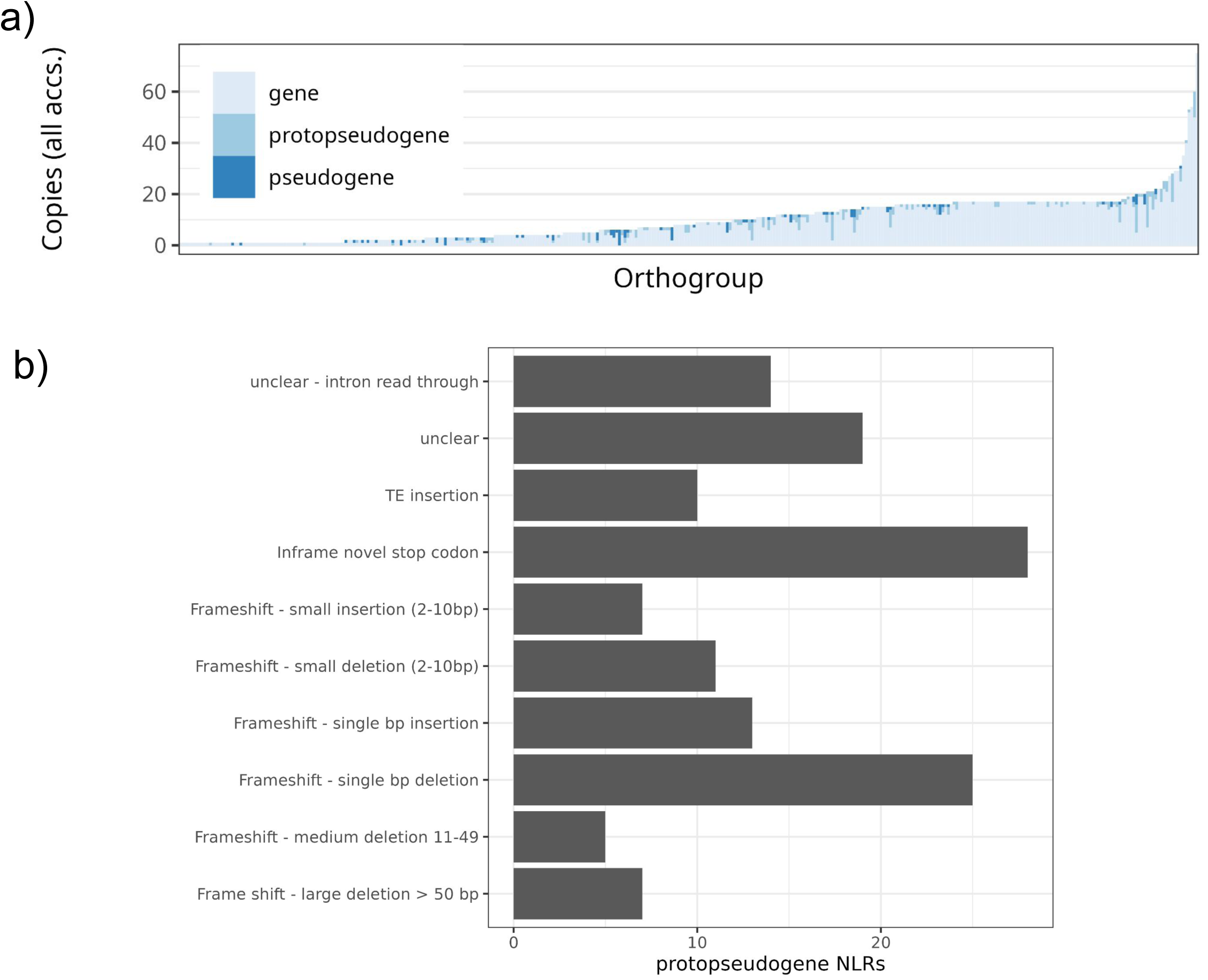
Pseudogenisation distribution and cause. a) the distribution of pseudogenes and protopseuodgenes across the fine orthogroups. b) The frequency of causes of protopseudogenisation/truncation in this dataset.

**Supplementary Figure 12:**
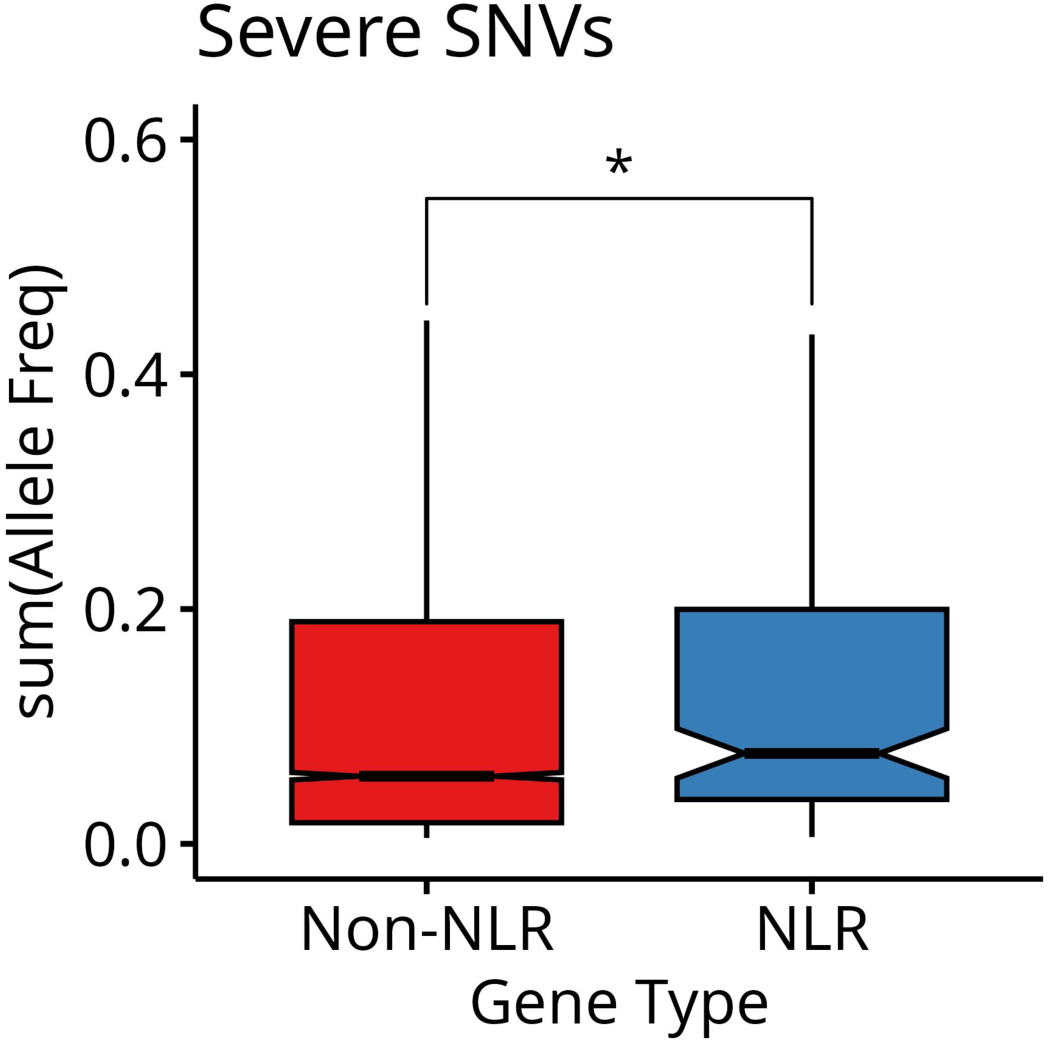
Severe mutations in 1135 short read genomes relative to at9852. NLRs have a significantly elevated rate of severe mutations compared to all Non-NLR genes.

**Supplementary Figure 14:**
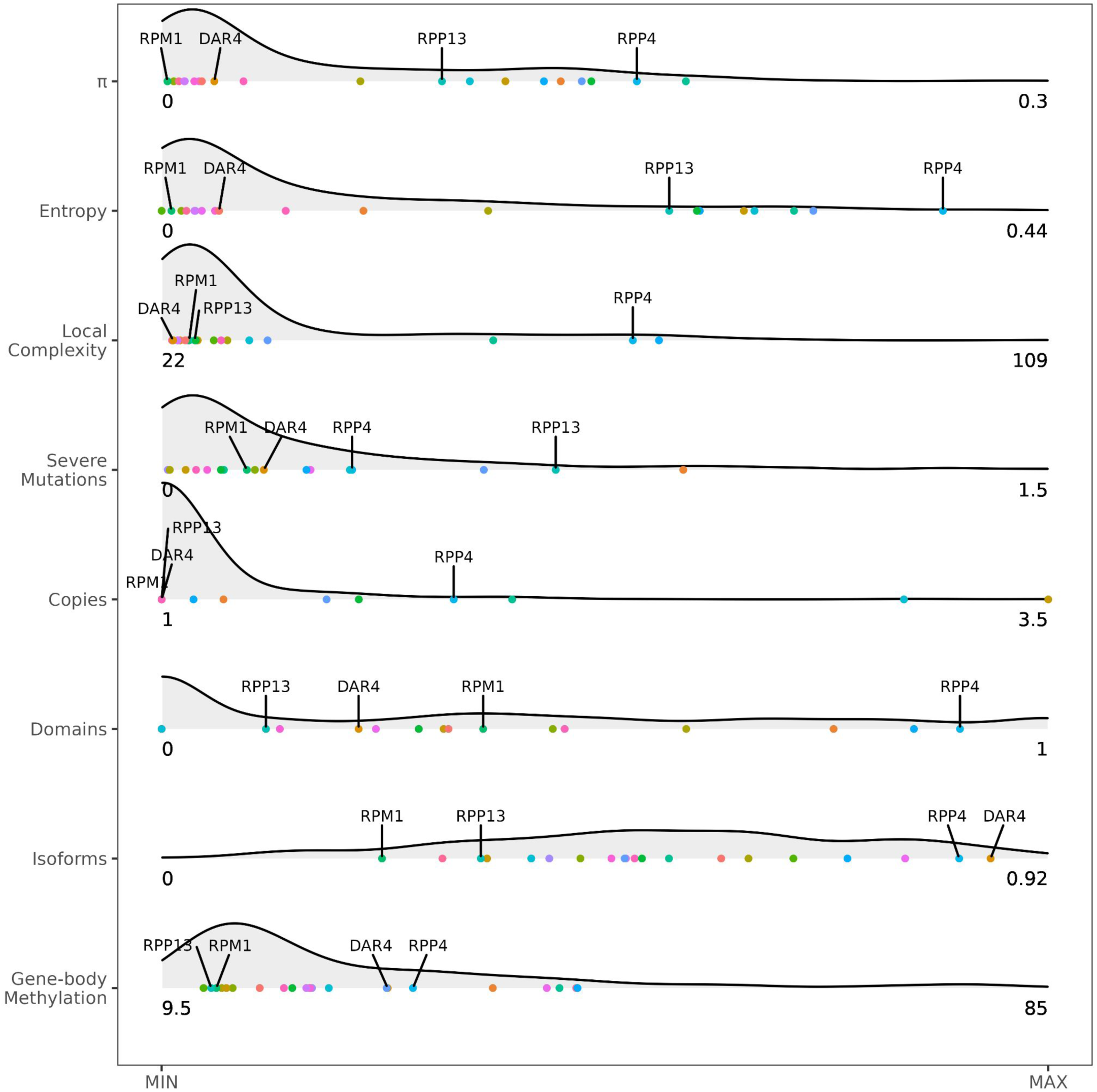
An enlarged representation of the distribution of known R genes along the axes of diversity considered here.

**Supplementary Figure 15:**
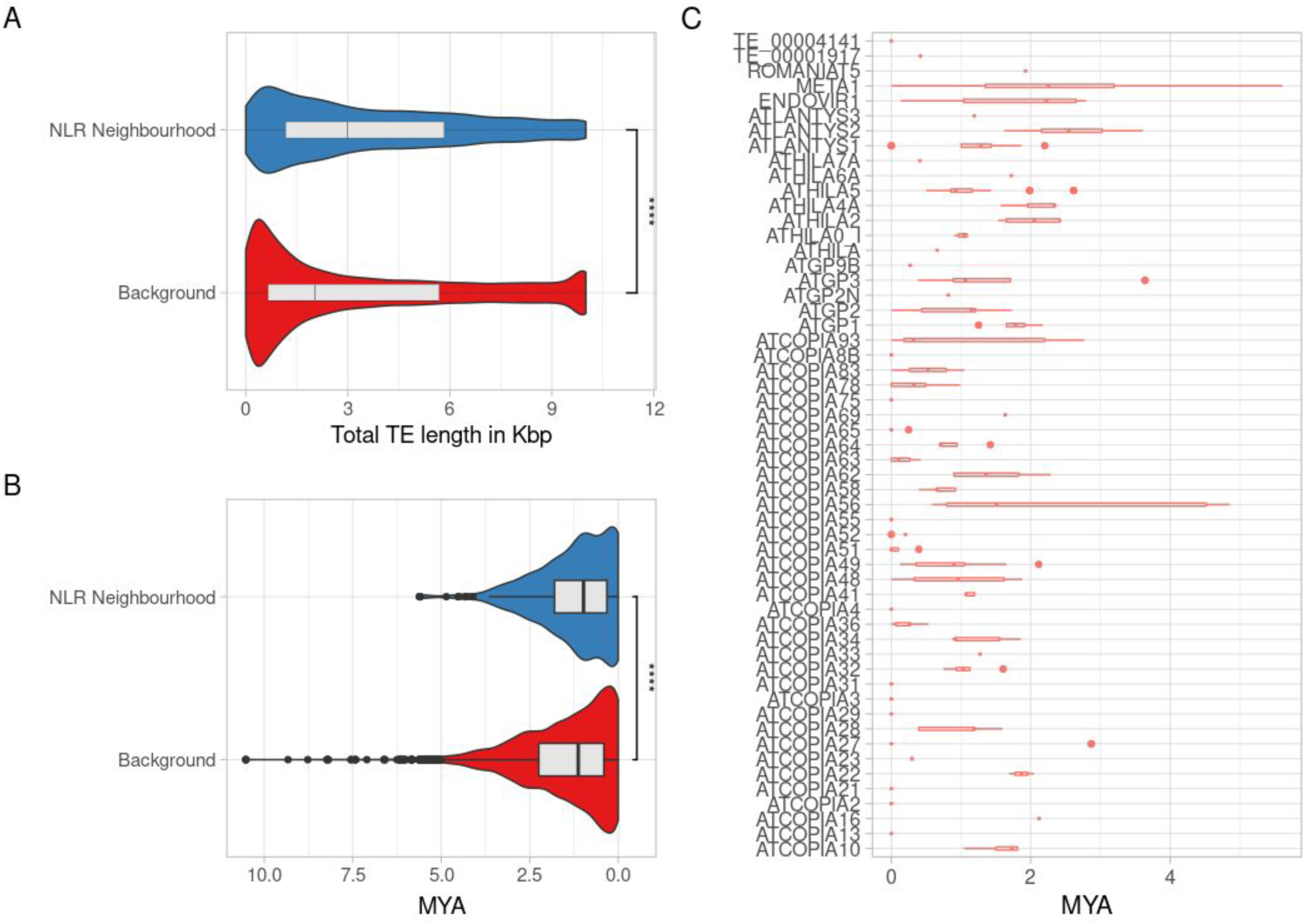
TEs within NLR neighborhoods: To compare transposable elements (TEs) within NLR neighborhoods to those in the genomic background, we first segmented the genomes into 10 Kbp windows, excluding telomeric, centromeric, and rRNA regions. These windows were then classified as either NLR neighborhood windows or background windows based on their overlap with NLR neighborhoods. After categorizing the windows, we performed the following analyses: A) calculated the TE density in each category, expressed as the total TE content per Kbp and compared using a t-test (p-value < 2.22e-16); and B) assessed the age of LTRs within each category (see methods) and compared using a t-test (p-value 8.2e-07). Panel C) shows the age variance of the different TE families present in the NLR neighborhoods.

**Supplementary Figure 16:**
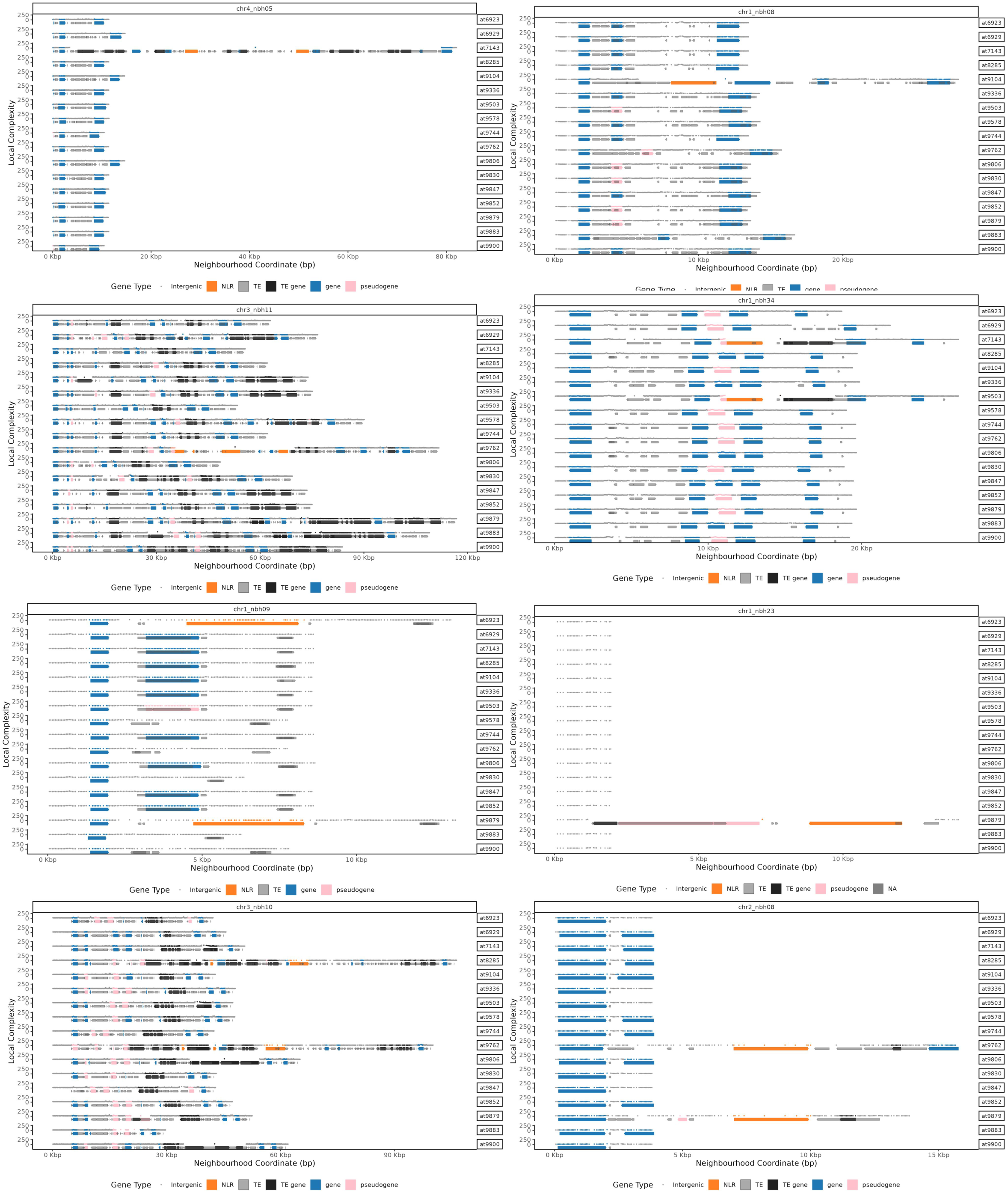
Probable cases of TE-mediated distal copying of NLRs. These neighbourhoods represent likely cases of TE-mediate copy-pasting of NLRs to distal locations. In all cases, NLRs appear in only a few accessions as part of a distinct chunk of DNA rich in TEs. In total, at least 12 such neighbourhoods occur.

**Supplementary Figure 17:**
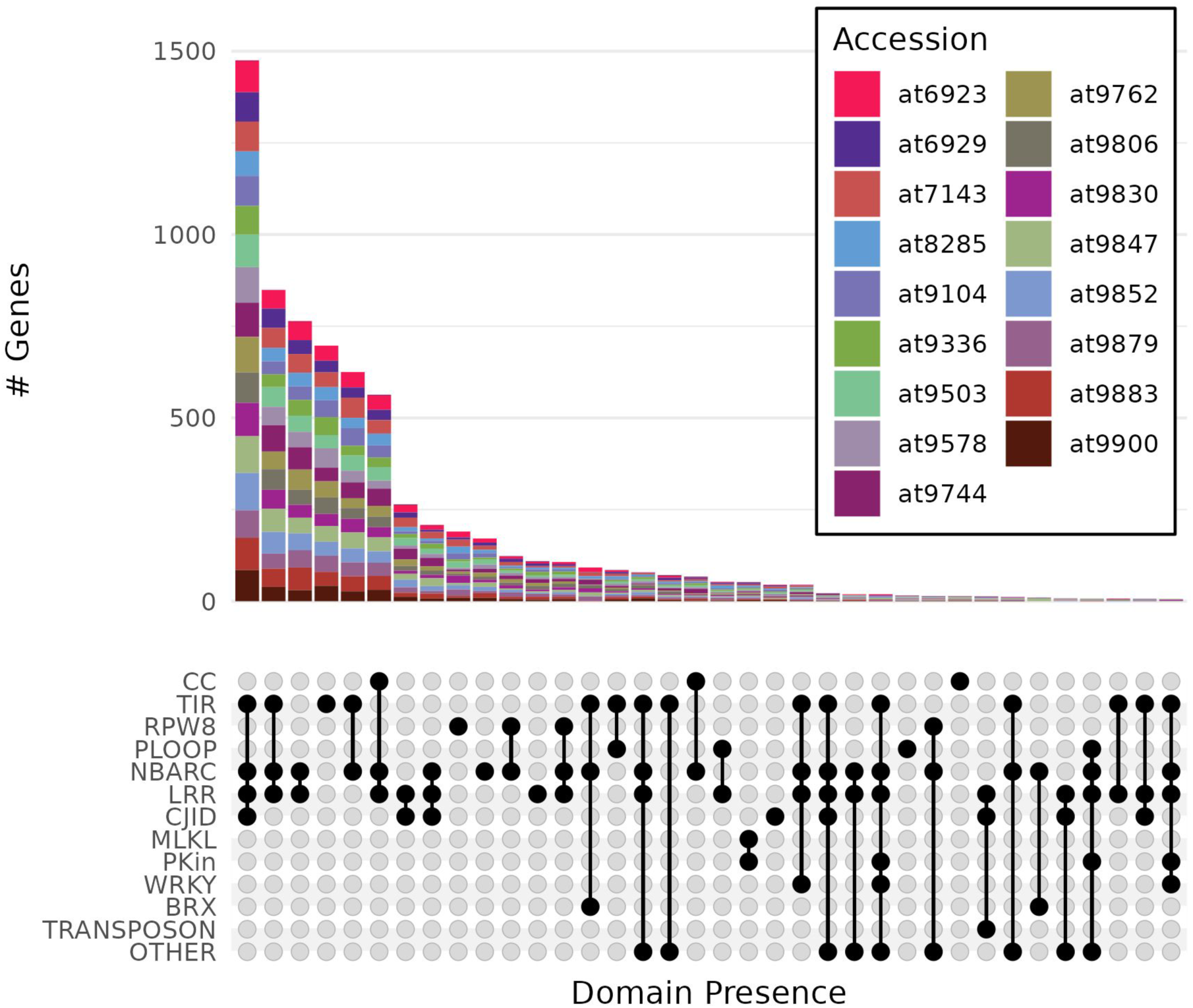
Domain presence absence in the NLRs across the 17 genomes.

**Supplementary Figure 18:**
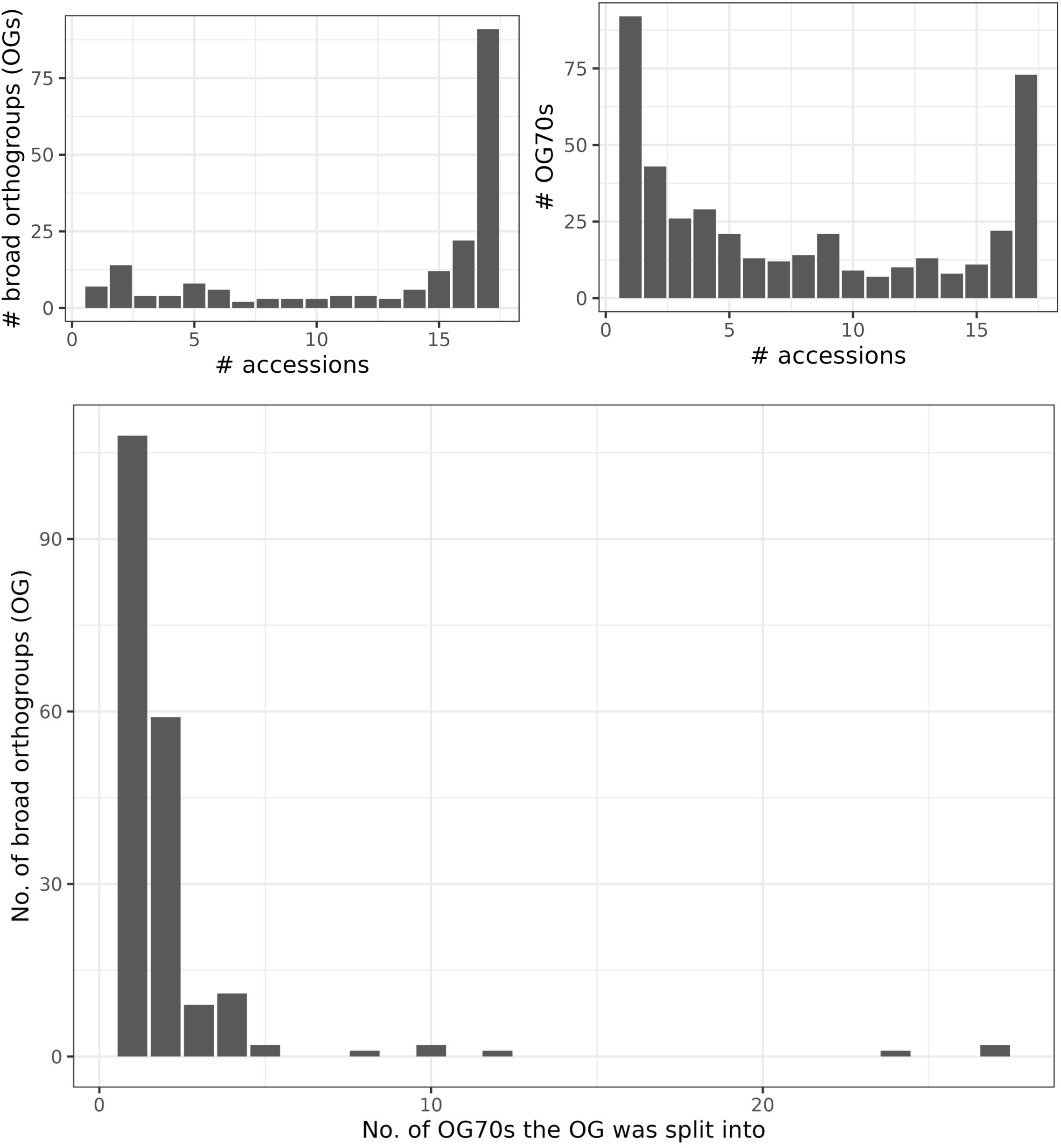
The distribution of the number of accessions with at least one copy of each orthogroup. Most NLR fine orthogroups are unique or present in every accession.

**Supplementary Figure 19:**
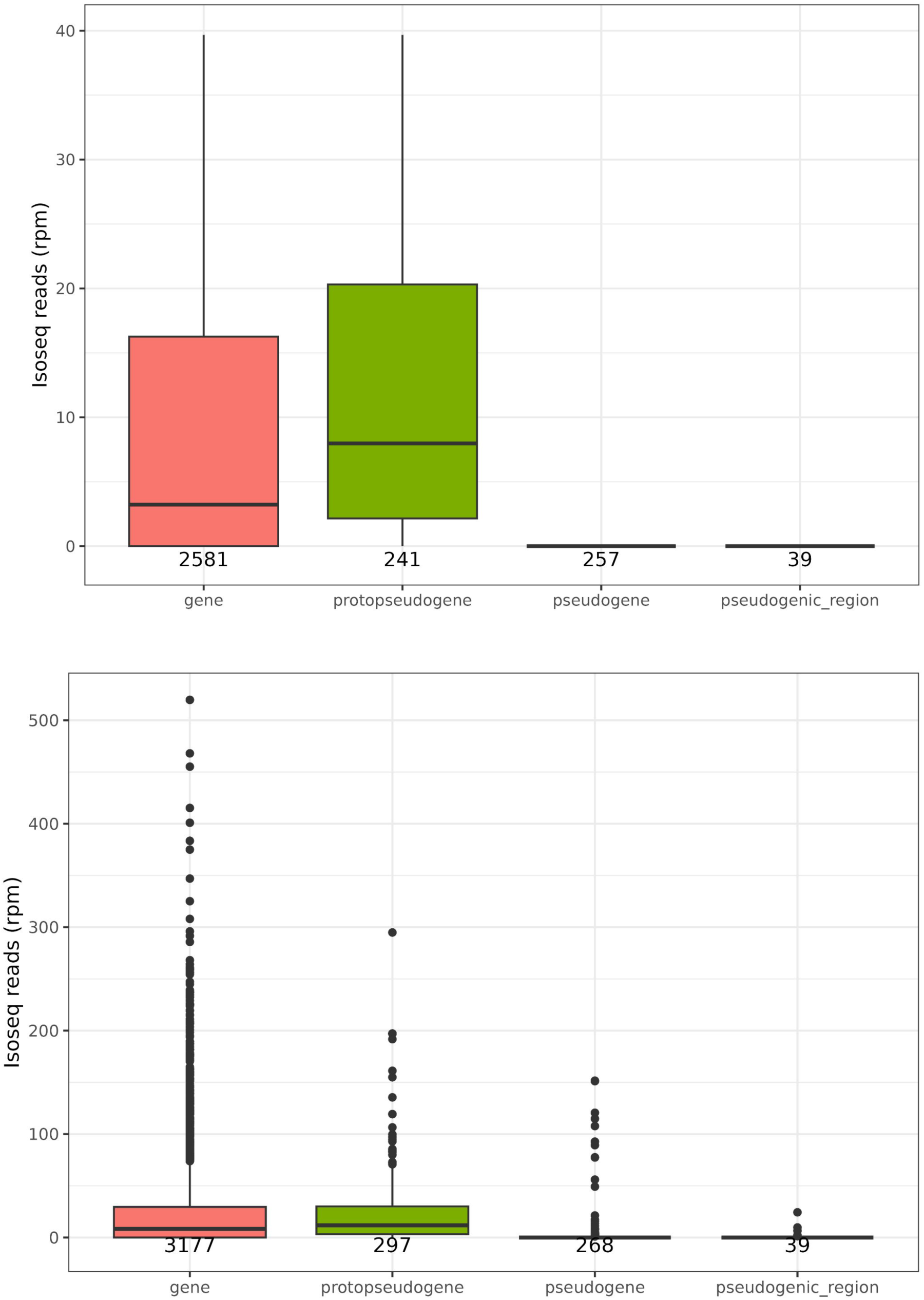
NLR genes and proto-pseudogenes are broadly expressed, to varying degrees, whereas very few pseudogenes and NLR remnants are expressed a) without outliers and b) is with outliers. All counts are in reads per million valid mapped reads.

**Supplementary Figure 20:**
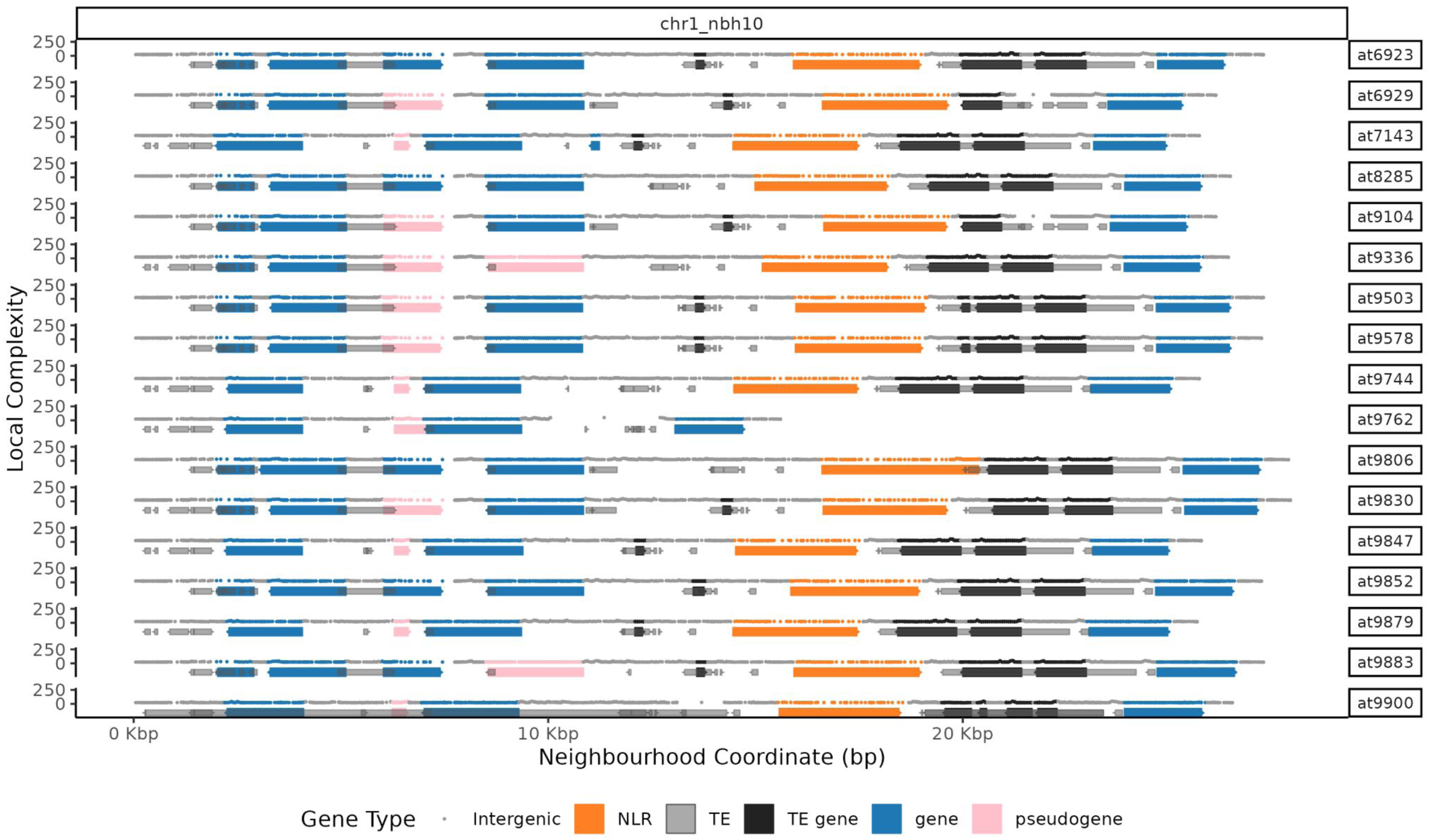
Probable case of TE-mediated deletion of ADR1. ADR1 is a highly-conserved and critical helper NLR. Nonetheless, we see one occurrence (in at9762) of what appears to be TE-mediated deletion of the ADR1-containing locus, including bordering TE genes and other TE sequence.

**Supplementary Figure 21:**
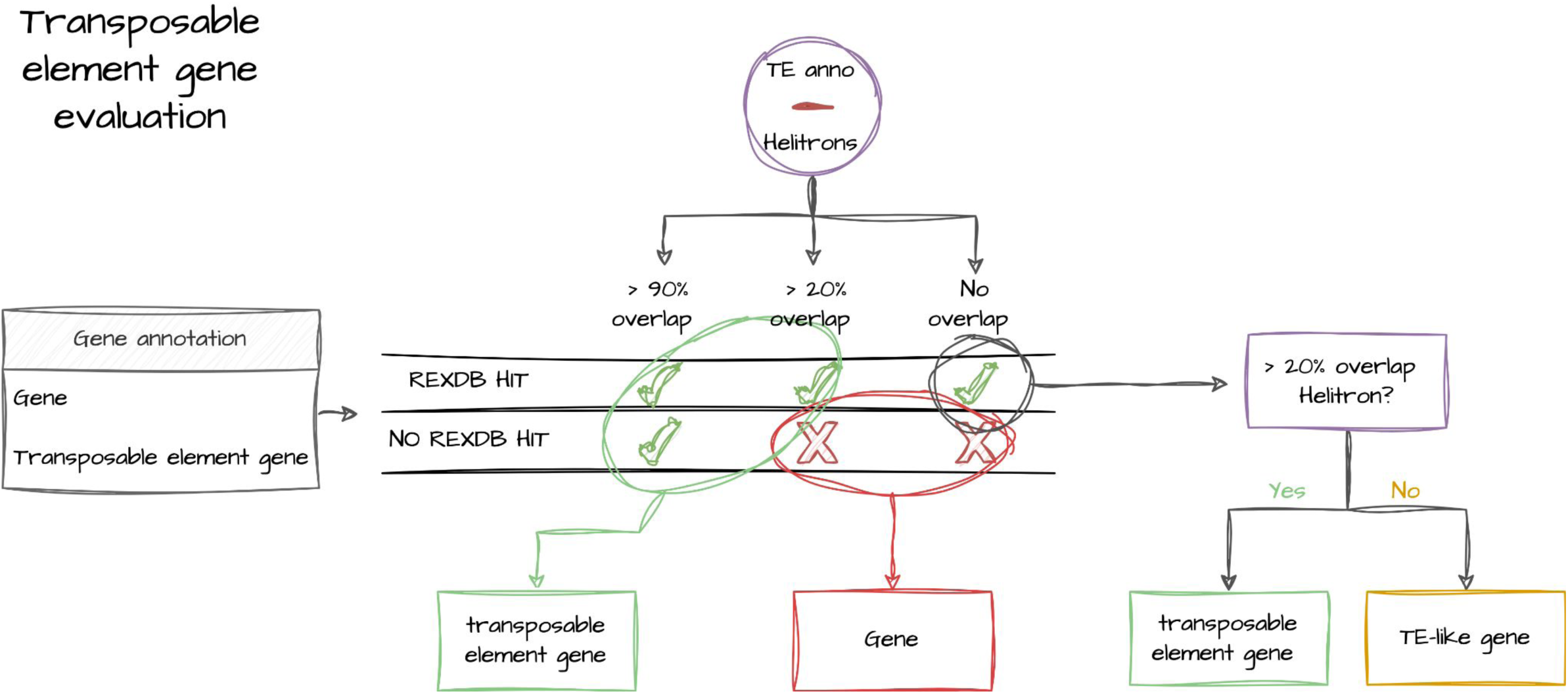
TE protein-coding gene annotation pipeline. We classified genes as transposable element genes if they had more than 90% of their annotated sequence overlapping with an annotated TE (other than helitron), or if they had an overlap between 20% and 90% but also contained in their sequence any TE protein domain present in the RexDB database. Genes with a RexDB hit and more than 20% overlap with helitrons were also classified as transposable element genes. The remaining genes that show a hit against RexDB and no overlap with any annotated TEs were classified as TE-like genes.

**Supplementary Figure 22:**
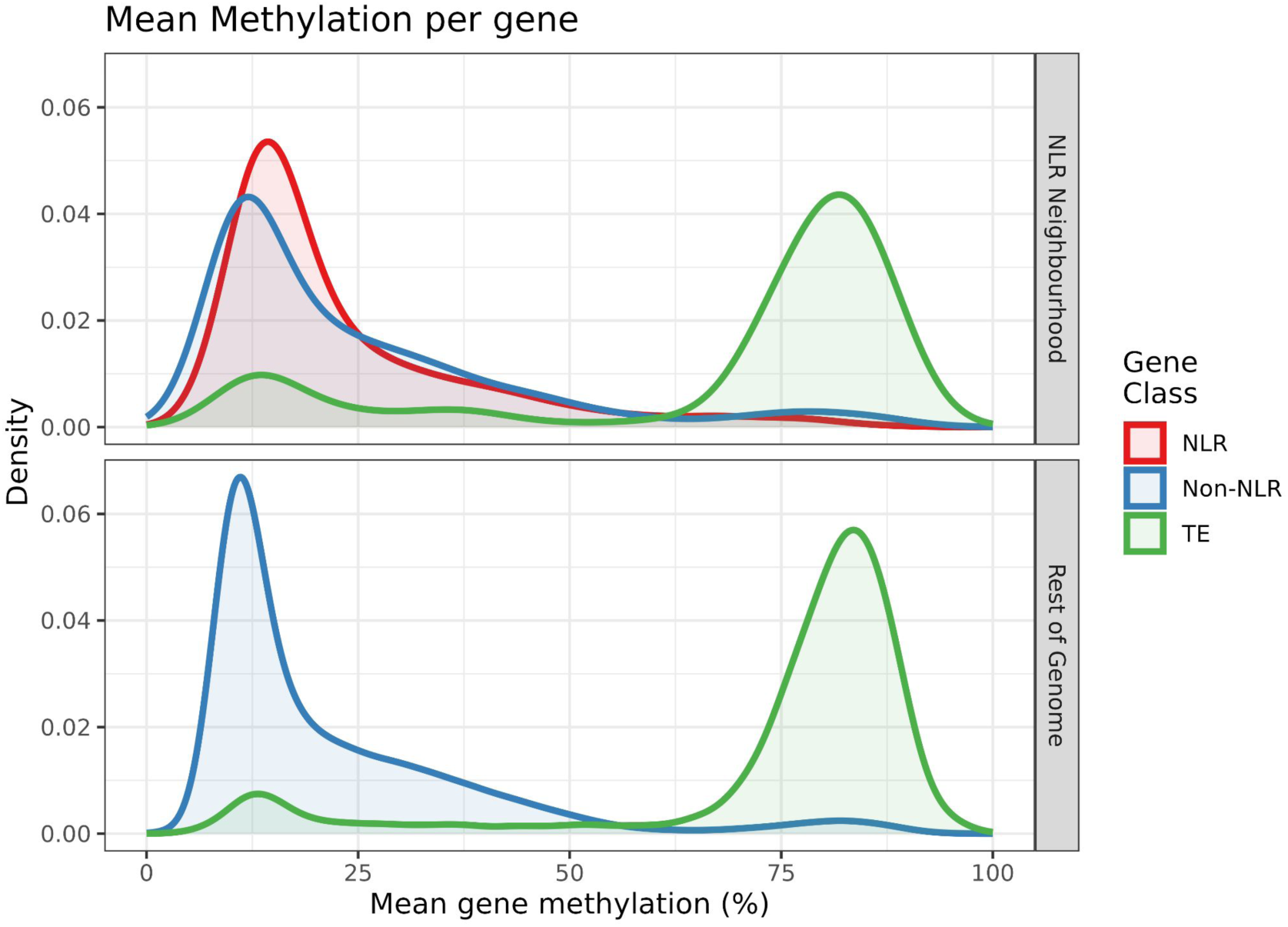
Steady state CG Methlylation across the 17 accessions. We show the distribution of the average steady state CG methylation per annotation (NLR gene, Non-NLR gene, and TE) in the NLR neighbourhoods compared to the rest of the genome (with repeat regions masked).

**Supplementary Figure 23:**
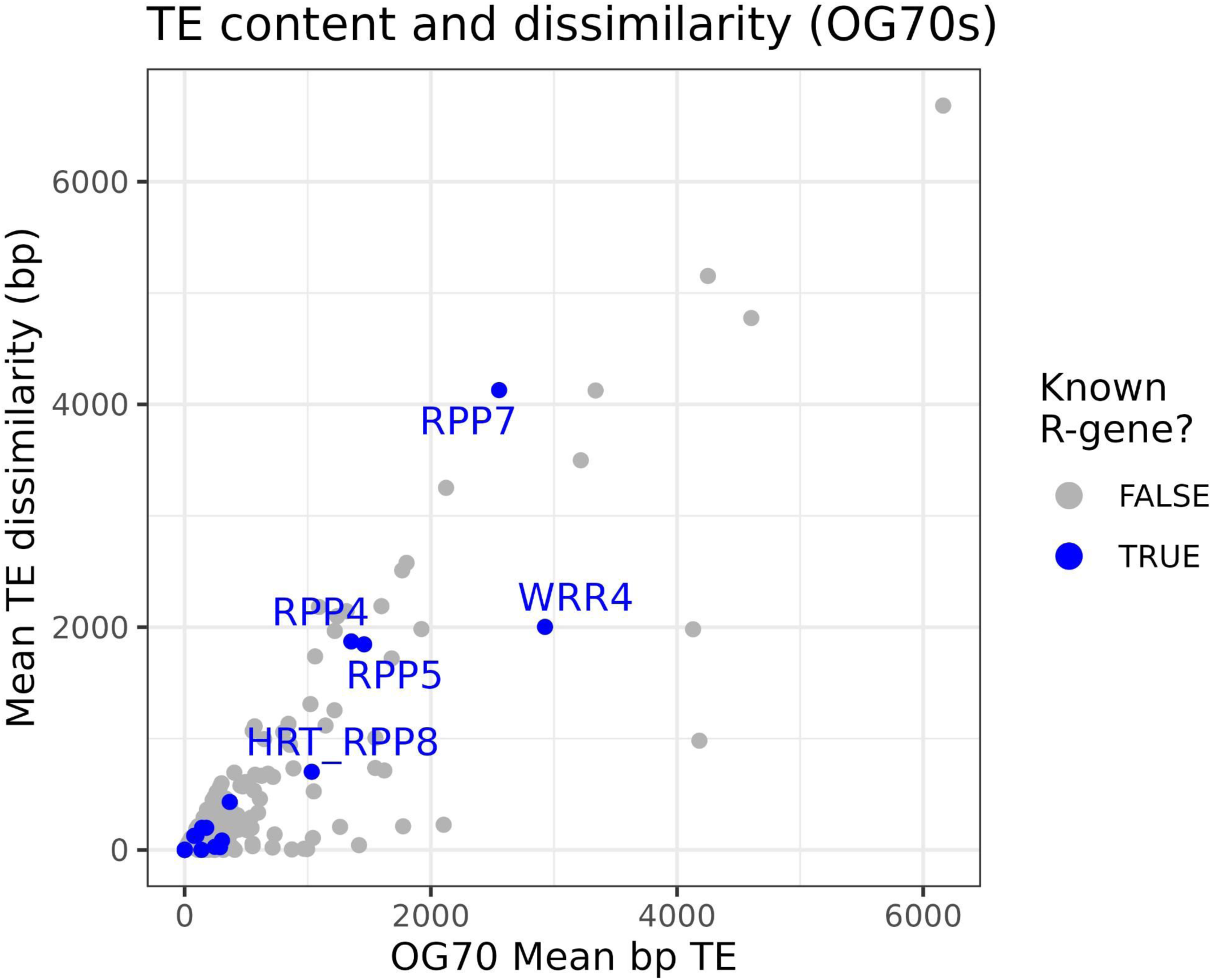
The relationship between the average *intragenic* TE content dissimilarity across all members of an OG70 and the average amount of TE present within the gene sequences belonging to the OG70. While many NLRs contain TEs, the orthogroups vary in the amount of TE variation. TE content dissimilarity counts the number of basepairs differentially annotated by TE family between two accessions. For example, if in some gene/region accession A has a total of 1000 bp each of TE families X and Y, while accession B has a total of 1000 bp each of TE families X and Z, while accession C has no TEs annotated, the distances among all accessions are 2000 bp. Below, we report the mean pairwise TE content dissimarlity between all OG70 members, and mean length of TE within the gene extent per OG70.

## Notes

https://dl20.coevolutionlab.org/datarelease/

### Competing Interest Statement

Detlef Weigel holds equity in Computomics, which advises plant breeders. Detlef Weigel also consults for KWS SE, a globally active plant breeder and seed producer. The other authors declare no competing interests.

## References

Alonge M, Lebeigle L, Kirsche M, Jenike K, Ou S, Aganezov S, Wang X, Lippman ZB, Schatz MC, Soyk S. 2022. Automated assembly scaffolding using RagTag elevates a new tomato system for high-throughput genome editing. Genome Biol. 23:258.

Alonso-Blanco C, Andrade J, Becker C, Bemm F, Bergelson J, Borgwardt KM, Cao J, Chae E, Dezwaan TM, Ding W, et al. 2016. 1,135 Genomes Reveal the Global Pattern of Polymorphism in Arabidopsis thaliana. Cell 166:481–491.

Ameline-Torregrosa C, Wang B-B, O’Bleness MS, Deshpande S, Zhu H, Roe B, Young ND, Cannon SB. 2007. Identification and Characterization of Nucleotide-Binding Site-Leucine-Rich Repeat Genes in the Model Plant Medicago truncatula. Plant Physiol. 146:5–21.

Bakker EG, Toomajian C, Kreitman M, Bergelson J. 2006. A Genome-Wide Survey of R Gene Polymorphisms in Arabidopsis. Plant Cell 18:1803–1818.

Barragan AC, Weigel D. 2021. Plant NLR diversity: the known unknowns of pan-NLRomes. Plant Cell 33:814–831.

Baumgarten A, Cannon S, Spangler R, May G. 2003. Genome-level evolution of resistance genes in Arabidopsis thaliana. Genetics 165:309–319.

Bertazzoni S, Williams AH, Jones DA, Syme RA, Tan K-C, Hane JK. 2018. Accessories make the outfit: Accessory chromosomes and other dispensable DNA regions in plant-pathogenic fungi. Mol. Plant. Microbe. Interact. 31:779–788.

van der Biezen EA, Freddie CT, Kahn K, Parker JE, Jones JDG. 2002. Arabidopsis RPP4 is a member of the RPP5 multigene family of TIR-NB-LRR genes and confers downy mildew resistance through multiple signalling components. Plant J. 29:439–451.

Bittner-Eddy PD, Crute IR, Holub EB, Beynon JL. 2000. RPP13 is a simple locus in Arabidopsis thaliana for alleles that specify downy mildew resistance to different avirulence determinants in Peronospora parasitica. Plant J. 21:177–188.

Boehm T, Swann JB. 2014. Origin and evolution of adaptive immunity. Annu. Rev. Anim. Biosci. 2:259–283.

Brabham HJ, Hernández-Pinzón I, Yanagihara C, Ishikawa N, Komori T, Matny ON, Hubbard A, Witek K, Feist A, Numazawa H, et al. 2024. Discovery of functional NLRs using expression level, high-throughput transformation, and large-scale phenotyping. bioRxiv [Internet]:2024.06.25.599845. Available from: https://www.biorxiv.org/content/10.1101/2024.06.25.599845v1

Brown JKM, Tellier A. 2011. Plant-parasite coevolution: bridging the gap between genetics and ecology. Annu. Rev. Phytopathol. 49:345–367.

Brůna T, Hoff KJ, Lomsadze A, Stanke M, Borodovsky M. 2021. BRAKER2: automatic eukaryotic genome annotation with GeneMark-EP+ and AUGUSTUS supported by a protein database. NAR Genom Bioinform 3:lqaa108.

Buchfink B, Reuter K, Drost H-G. 2021. Sensitive protein alignments at tree-of-life scale using DIAMOND. Nat. Methods 18:366–368.

Camacho C, Coulouris G, Avagyan V, Ma N, Papadopoulos J, Bealer K, Madden TL. 2009. BLAST+: architecture and applications. BMC Bioinformatics 10:421.

Chae E, Bomblies K, Kim S-T, Karelina D, Zaidem M, Ossowski S, Martín-Pizarro C, Laitinen RAE, Rowan BA, Tenenboim H, et al. 2014. Species-wide genetic incompatibility analysis identifies immune genes as hot spots of deleterious epistasis. Cell 159:1341–1351.

Cheetham SW, Kindlova M, Ewing AD. 2022. Methylartist: tools for visualizing modified bases from nanopore sequence data. Bioinformatics 38:3109–3112.

Cheng C-Y, Krishnakumar V, Chan AP, Thibaud-Nissen F, Schobel S, Town CD. 2017. Araport11: a complete reannotation of the Arabidopsis thaliana reference genome. Plant J. 89:789–804.

Cheng H, Concepcion GT, Feng X, Zhang H, Li H. 2021. Haplotype-Resolved de Novo Assembly Using Phased Assembly Graphs with Hifiasm. Nat. Methods 18:170–175.

Chen N. 2004. Using RepeatMasker to identify repetitive elements in genomic sequences. Curr. Protoc. Bioinformatics Chapter 4:Unit 4.10.

Choi K, Reinhard C, Serra H, Ziolkowski PA, Underwood CJ, Zhao X, Hardcastle TJ, Yelina NE, Griffin C, Jackson M, et al. 2016. Recombination Rate Heterogeneity within Arabidopsis Disease Resistance Genes. PLoS Genet. 12:e1006179.

Clark RM, Schweikert G, Toomajian C, Ossowski S, Zeller G, Shinn P, Warthmann N, Hu TT, Fu G, Hinds DA, et al. 2007. Common sequence polymorphisms shaping genetic diversity in Arabidopsis thaliana. Science 317:338–342.

Dainat J, Hereñú D, Davis E, Crouch K, LucileSol, Agostinho N, pascal-git, Zollman Z, tayyrov. 2023. NBISweden/AGAT: AGAT-v1.1.0. Zenodo Available from: https://zenodo.org/record/3552717

Danecek P, Bonfield JK, Liddle J, Marshall J, Ohan V, Pollard MO, Whitwham A, Keane T, McCarthy SA, Davies RM, et al. 2021. Twelve years of SAMtools and BCFtools. Gigascience 10:1–4.

Danecek P, McCarthy SA. 2017. BCFtools/csq: haplotype-aware variant consequences. Bioinformatics 33:2037–2039.

Dodds PN, Rathjen JP. 2010. Plant Immunity: Towards an Integrated View of Plant-Pathogen Interactions. Nat. Rev. Genet. 11:539–548.

Ellinghaus D, Kurtz S, Willhoeft U. 2008. LTRharvest, an efficient and flexible software for de novo detection of LTR retrotransposons. BMC Bioinformatics 9:18.

Emms DM, Kelly S. 2019. OrthoFinder: phylogenetic orthology inference for comparative genomics. Genome Biol. 20:238.

Gan X, Stegle O, Behr J, Steffen JG, Drewe P, Hildebrand KL, Lyngsoe R, Schultheiss SJ, Osborne EJ, Sreedharan VT, et al. 2011. Multiple reference genomes and transcriptomes for Arabidopsis thaliana. Nature 477:419–423.

Garrison E, Guarracino A, Heumos S, Villani F, Bao Z, Tattini L, Hagmann J, Vorbrugg S, Marco-Sola S, Kubica C, et al. 2023. Building pangenome graphs. bioRxiv [Internet]. Available from: 10.1101/2023.04.05.535718

Garrison E, Marth G. 2012. Haplotype-Based Variant Detection from Short-Read Sequencing. arXiv:1207. 3907 [q-bio] [Internet]. Available from: http://arxiv.org/abs/1207.3907

Giorgetti OB, O’Meara CP, Schorpp M, Boehm T. 2023. Origin and evolutionary malleability of T cell receptor α diversity. Nature 619:193–200.

Goel M, Sun H, Jiao W-B, Schneeberger K. 2019. SyRI: finding genomic rearrangements and local sequence differences from whole-genome assemblies. Genome Biol. 20:277.

Guarracino A, Heumos S, Nahnsen S, Prins P, Garrison E. 2022. ODGI: understanding pangenome graphs. Bioinformatics 38:3319–3326.

Gurevich A, Saveliev V, Vyahhi N, Tesler G. 2013. QUAST: quality assessment tool for genome assemblies. Bioinformatics 29:1072–1075.

Haas BJ, Delcher AL, Mount SM, Wortman JR, Smith RK Jr, Hannick LI, Maiti R, Ronning CM, Rusch DB, Town CD, et al. 2003. Improving the Arabidopsis genome annotation using maximal transcript alignment assemblies. Nucleic Acids Res. 31:5654–5666.

Haas B, Papanicolaou A, Yassour M. 2017. TransDecoder. Available from: https://github.com/TransDecoder/TransDecoder

Han Y, Wessler SR. 2010. MITE-Hunter: a program for discovering miniature inverted-repeat transposable elements from genomic sequences. Nucleic Acids Res. 38:e199.

Holub EB. 2001. The arms race is ancient history in Arabidopsis, the wildflower. Nat. Rev. Genet. 2:516–527.

Hu K, Xu K, Wen J, Yi B, Shen J, Ma C, Fu T, Ouyang Y, Tu J. 2019. Helitron distribution in Brassicaceae and whole Genome Helitron density as a character for distinguishing plant species. BMC Bioinformatics 20:354.

Jiao W-B, Schneeberger K. 2020. Chromosome-level assemblies of multiple Arabidopsis genomes reveal hotspots of rearrangements with altered evolutionary dynamics. Nat. Commun. 11:1–10.

Jones JDG, Staskawicz BJ, Dangl JL. 2024. The plant immune system: From discovery to deployment. Cell 187:2095–2116.

Jones P, Binns D, Chang H-Y, Fraser M, Li W, McAnulla C, McWilliam H, Maslen J, Mitchell A, Nuka G, et al. 2014. InterProScan 5: genome-scale protein function classification. Bioinformatics 30:1236–1240.

Karasov TL, Chae E, Herman JJ, Bergelson J. 2017. Mechanisms to Mitigate the Trade-Off between Growth and Defense. Plant Cell 29:666–680.

Katoh K, Kuma K-I, Toh H, Miyata T. 2005. MAFFT Version 5: Improvement in Accuracy of Multiple Sequence Alignment. Nucleic Acids Res. 33:511–518.

Kawakatsu T, Huang S-SC, Jupe F, Sasaki E, Schmitz RJ, Urich MA, Castanon R, Nery JR, Barragan C, He Y, et al. 2016. Epigenomic Diversity in a Global Collection of Arabidopsis Thaliana Accessions. Cell 166:492–505.

Koenig D, Hagmann J, Li R, Bemm F, Slotte T, Neuffer B, Wright SI, Weigel D. 2019. Long-term balancing selection drives evolution of immunity genes in Capsella. Elife 8:e43606.

Kourelis J, Sakai T, Adachi H, Kamoun S. 2021. RefPlantNLR is a comprehensive collection of experimentally validated plant disease resistance proteins from the NLR family. PLoS Biol. 19:e3001124.

Krueger F, Andrews SR. 2011. Bismark: a flexible aligner and methylation caller for Bisulfite-Seq applications. Bioinformatics 27:1571–1572.

Kufel J, Diachenko N, Golisz A. 2022. Alternative splicing as a key player in the fine-tuning of the immunity response in Arabidopsis. Mol. Plant Pathol. 23:1226–1238.

Kuo RI, Cheng Y, Zhang R, Brown JWS, Smith J, Archibald AL, Burt DW. 2020. Illuminating the dark side of the human transcriptome with long read transcript sequencing. BMC Genomics 21:751.

Laetsch DR, Blaxter ML. 2017. BlobTools: Interrogation of genome assemblies. F1000Res. 6:1287.

Lee E, Helt GA, Reese JT, Munoz-Torres MC, Childers CP, Buels RM, Stein L, Holmes IH, Elsik CG, Lewis SE. 2013. Web Apollo: a web-based genomic annotation editing platform. Genome Biol. 14:R93.

Lee RRQ, Chae E. 2020. Patterns of NLR cluster variation in Arabidopsis thaliana genomes. Plant Communications 1:100089.

Liang W, van Wersch S, Tong M, Li X. 2019. TIR-NB-LRR immune receptor SOC3 pairs with truncated TIR-NB protein CHS1 or TN2 to monitor the homeostasis of E3 ligase SAUL1. New Phytol. 221:2054–2066.

Lian Q, Huettel B, Walkemeier B, Mayjonade B, Lopez-Roques C, Gil L, Roux F, Schneeberger K, Mercier R. 2024. A pan-genome of 69 Arabidopsis thaliana accessions reveals a conserved genome structure throughout the global species range. Nat. Genet. 56:982–991.

Li H. 2008. Seqtk - Toolkit for Processing Sequences in FASTA/Q Formats. Available from: https://github.com/lh3/seqtk

Li H. 2013. Aligning Sequence Reads, Clone Sequences and Assembly Contigs with BWA-MEM. arXiv [Internet]. Available from: 10.48550/arXiv.1303.3997

Li H. 2018. Minimap2: pairwise alignment for nucleotide sequences. Bioinformatics 34:3094–3100.

Li H. 2021. New strategies to improve minimap2 alignment accuracy. Bioinformatics 37:4572–4574.

Li H, Handsaker B, Wysoker A, Fennell T, Ruan J, Homer N, Marth G, Abecasis G, Durbin R, 1000 Genome Project Data Processing Subgroup. 2009. The Sequence Alignment/Map format and SAMtools. Bioinformatics 25:2078–2079.

Li W, Godzik A. 2006. Cd-hit: a fast program for clustering and comparing large sets of protein or nucleotide sequences. Bioinformatics 22:1658–1659.

Lovell JT, Sreedasyam A, Schranz ME, Wilson M, Carlson JW, Harkess A, Emms D, Goodstein DM, Schmutz J. 2022. GENESPACE tracks regions of interest and gene copy number variation across multiple genomes. Elife 11:e78526.

MacQueen A, Bergelson J. 2016. Modulation of R-gene expression across environments. J. Exp. Bot. 67:2093–2105.

Mencia R, Arce AL, Houriet C, Xian W, Contreras A, Shirsekar G, Weigel D, Manavella PA. 2023. Transposon-triggered epigenetic chromatin dynamics modulate EFR-related pathogen response. bioRxiv [Internet]:2023.10.06.561201. Available from: https://www.biorxiv.org/content/10.1101/2023.10.06.561201v1

Michelmore RW, Meyers BC. 1998. Clusters of resistance genes in plants evolve by divergent selection and a birth-and-death process. Genome Res. 8:1113–1130.

Minh BQ, Schmidt HA, Chernomor O, Schrempf D, Woodhams MD, von Haeseler A, Lanfear R. 2020. IQ-TREE 2: New Models and Efficient Methods for Phylogenetic Inference in the Genomic Era. Mol. Biol. Evol. 37:1530–1534.

Möller M, Stukenbrock EH. 2017. Evolution and genome architecture in fungal plant pathogens. Nat. Rev. Microbiol. 15:756–771.

Monroe JG, McKay JK, Weigel D, Flood PJ. 2021. The population genomics of adaptive loss of function. Heredity 126:383–395.

Monroe JG, Powell T, Price N, Mullen JL, Howard A, Evans K, Lovell JT, McKay JK. 2018. Drought adaptation in Arabidopsis thaliana by extensive genetic loss-of-function. Elife 7:e41038.

Murray KD. 2024a. kdm9/blindschleiche: Version 0.3.1. Zenodo Available from: https://zenodo.org/doi/10.5281/zenodo.10049825

Murray KD. 2024b. kdm9/raugraf: Version 0.0.5. Zenodo Available from: https://zenodo.org/doi/10.5281/zenodo.13144148

Murray KD, Borevitz JO, Weigel D, Warthmann N. 2024. Acanthophis: a comprehensive plant hologenomics pipeline. J. Open Source Softw. 9:6062.

Nachtweide S, Stanke M. 2019. Multi-Genome Annotation with AUGUSTUS. Methods Mol. Biol. 1962:139–160.

Nemri A, Atwell S, Tarone AM, Huang YS, Zhao K, Studholme DJ, Nordborg M, Jones JDG. 2010. Genome-wide survey of Arabidopsis natural variation in downy mildew resistance using combined association and linkage mapping. Proc. Natl. Acad. Sci. U. S. A. 107:10302–10307.

Neumann P, Novák P, Hoštáková N, Macas J. 2019. Systematic survey of plant LTR-retrotransposons elucidates phylogenetic relationships of their polyprotein domains and provides a reference for element classification. Mob. DNA 10:1.

Ni P, Nie F, Zhong Z, Xu J, Huang N, Zhang J, Zhao H, Zou Y, Huang Y, Li J, et al. 2023. DNA 5-methylcytosine detection and methylation phasing using PacBio circular consensus sequencing. Nat. Commun. 14:4054.

Nishimura MT, Anderson RG, Cherkis KA, Law TF, Liu QL, Machius M, Nimchuk ZL, Yang L, Chung E-H, El Kasmi F, et al. 2017. TIR-only protein RBA1 recognizes a pathogen effector to regulate cell death in Arabidopsis. Proc. Natl. Acad. Sci. U. S. A. 114:E2053–E2062.

Ossowski S, Schneeberger K, Lucas-Lledó JI, Warthmann N, Clark RM, Shaw RG, Weigel D, Lynch M. 2010. The rate and molecular spectrum of spontaneous mutations in Arabidopsis thaliana. Science 327:92–94.

Ou S, Jiang N. 2018. LTR_retriever: A Highly Accurate and Sensitive Program for Identification of Long Terminal Repeat Retrotransposons. Plant Physiol. 176:1410–1422.

Ou S, Jiang N. 2019. LTR_FINDER_parallel: parallelization of LTR_FINDER enabling rapid identification of long terminal repeat retrotransposons. Mob. DNA 10:48.

Ou S, Su W, Liao Y, Chougule K, Agda JRA, Hellinga AJ, Lugo CSB, Elliott TA, Ware D, Peterson T, et al. 2019. Benchmarking transposable element annotation methods for creation of a streamlined, comprehensive pipeline. Genome Biol. 20:275.

Panda K, Slotkin RK. 2020. Long-Read cDNA Sequencing Enables a “Gene-Like” Transcript Annotation of Transposable Elements. Plant Cell 32:2687–2698.

Paradis E, Schliep K. 2019. ape 5.0: an environment for modern phylogenetics and evolutionary analyses in R. Bioinformatics 35:526–528.

Pardo-Palacios FJ, Arzalluz-Luque A, Kondratova L, Salguero P, Mestre-Tomás J, Amorín R, Estevan-Morió E, Liu T, Nanni A, McIntyre L, et al. 2024. SQANTI3: curation of long-read transcriptomes for accurate identification of known and novel isoforms. Nat. Methods 21:793–797.

Prigozhin DM, Krasileva KV. 2021. Analysis of intraspecies diversity reveals a subset of highly variable plant immune receptors and predicts their binding sites. Plant Cell 33:998–1015.

Rabanal FA, Gräff M, Lanz C, Fritschi K, Llaca V, Lang M, Carbonell-Bejerano P, Henderson I, Weigel D. 2022. Pushing the limits of HiFi assemblies reveals centromere diversity between two Arabidopsis thaliana genomes. Nucleic Acids Res. 50:12309–12327.

Reams AB, Roth JR. 2015. Mechanisms of gene duplication and amplification. Cold Spring Harb. Perspect. Biol. 7:a016592.

Robinson JT, Thorvaldsdottir H, Turner D, Mesirov JP. 2023. igv.js: an embeddable JavaScript implementation of the Integrative Genomics Viewer (IGV). Bioinformatics 39:btac830.

SanMiguel P, Gaut BS, Tikhonov A, Nakajima Y, Bennetzen JL. 1998. The paleontology of intergene retrotransposons of maize. Nat. Genet. 20:43–45.

Saucet SB, Ma Y, Sarris PF, Furzer OJ, Sohn KH, Jones JDG. 2015. Two linked pairs of Arabidopsis TNL resistance genes independently confer recognition of bacterial effector AvrRps4. Nat. Commun. 6:6338.

Schubert M, Lindgreen S, Orlando L. 2016. AdapterRemoval v2: rapid adapter trimming, identification, and read merging. BMC Res. Notes 9:88.

Shirsekar G, Devos J, Latorre SM, Blaha A, Queiroz Dias M, González Hernando A, Lundberg DS, Burbano HA, Fenster CB, Weigel D. 2021. Multiple Sources of Introduction of North American Arabidopsis thaliana from across Eurasia. Mol. Biol. Evol. 38:5328–5344.

Shumate A, Salzberg SL. 2021. Liftoff: Accurate Mapping of Gene Annotations. Bioinformatics 37:1639–1643.

Simão FA, Waterhouse RM, Ioannidis P, Kriventseva EV, Zdobnov EM. 2015. BUSCO: assessing genome assembly and annotation completeness with single-copy orthologs. Bioinformatics 31:3210–3212.

Steuernagel B, Witek K, Krattinger SG, Ramirez-Gonzalez RH, Schoonbeek H-J, Yu G, Baggs E, Witek AI, Yadav I, Krasileva KV, et al. 2020. The NLR-Annotator Tool Enables Annotation of the Intracellular Immune Receptor Repertoire. Plant Physiol. 183:468–482.

Stirnweis D, Milani SD, Brunner S, Herren G, Buchmann G, Peditto D, Jordan T, Keller B. 2014. Suppression among alleles encoding nucleotide-binding-leucine-rich repeat resistance proteins interferes with resistance in F1 hybrid and allele-pyramided wheat plants. Plant J. 79:893–903.

Sutherland CA, Prigozhin DM, Monroe JG, Krasileva KV. 2024. High allelic diversity in Arabidopsis NLRs is associated with distinct genomic features. EMBO Rep. 25:2306–2322.

Su W, Gu X, Peterson T. 2019. TIR-Learner, a New Ensemble Method for TIR Transposable Element Annotation, Provides Evidence for Abundant New Transposable Elements in the Maize Genome. Mol. Plant 12:447–460.

The Arabidopsis Genome Initiative. 2000. Analysis of the Genome Sequence of the Flowering Plant Arabidopsis Thaliana. Nature 408:796–815.

Thompson JN. 2005. The Geographic Mosaic of Coevolution. University of Chicago Press

Thrall PH, Burdon JJ. 2003. Evolution of virulence in a plant host-pathogen metapopulation. Science 299:1735–1737.

Tsuchiya T, Eulgem T. 2013. An alternative polyadenylation mechanism coopted to the Arabidopsis RPP7 gene through intronic retrotransposon domestication. Proc. Natl. Acad. Sci. U. S. A. 110:E3535–E3543.

Van de Weyer A-L, Monteiro F, Furzer OJ, Nishimura MT, Cevik V, Witek K, Jones JDG, Dangl JL, Weigel D, Bemm F. 2019. A Species-Wide Inventory of NLR Genes and Alleles in Arabidopsis thaliana. Cell 178:1260–1272.e14.

van Wersch S, Li X. 2019. Stronger When Together: Clustering of Plant NLR Disease resistance Genes. Trends Plant Sci. 24:688–699.

Xiong W, He L, Lai J, Dooner HK, Du C. 2014. HelitronScanner uncovers a large overlooked cache of Helitron transposons in many plant genomes. Proc. Natl. Acad. Sci. U. S. A. 111:10263–10268.

Yaffe H, Buxdorf K, Shapira I, Ein-Gedi S, Moyal-Ben Zvi M, Fridman E, Moshelion M, Levy M. 2012. LogSpin: a simple, economical and fast method for RNA isolation from infected or healthy plants and other eukaryotic tissues. BMC Res. Notes 5:45.

Yang S, Tang F, Zhu H. 2014. Alternative splicing in plant immunity. Int. J. Mol. Sci. 15:10424–10445.

Yang Y, Kim NH, Cevik V, Jacob P, Wan L, Furzer OJ, Dangl JL. 2022. Allelic variation in the Arabidopsis TNL CHS3/CSA1 immune receptor pair reveals two functional cell-death regulatory modes. Cell Host Microbe 30:1701–1716.e5.

Yi H, Richards EJ. 2007. A cluster of disease resistance genes in Arabidopsis is coordinately regulated by transcriptional activation and RNA silencing. Plant Cell 19:2929–2939.

Yuan W, Beitel F, Srikant T, Bezrukov I, Schäfer S, Kraft R, Weigel D. 2023. Pervasive under-dominance in gene expression underlying emergent growth trajectories in Arabidopsis thaliana hybrids. Genome Biol. 24:200.

Yu G. 2020. Using ggtree to Visualize Data on Tree-Like Structures. Curr. Protoc. Bioinformatics 69:e96.

Zdobnov EM, Kuznetsov D, Tegenfeldt F, Manni M, Berkeley M, Kriventseva EV. 2021. OrthoDB in 2020: evolutionary and functional annotations of orthologs. Nucleic Acids Res. 49:D389–D393.

Zervudacki J, Yu A, Amesefe D, Wang J, Drouaud J, Navarro L, Deleris A. 2018. Transcriptional control and exploitation of an immune-responsive family of plant retrotransposons. EMBO J. 37:e98482.

Zhang R-G, Li G-Y, Wang X-L, Dainat J, Wang Z-X, Ou S, Ma Y. 2022. TEsorter: an accurate and fast method to classify LTR-retrotransposons in plant genomes. Hortic Res 9:uhac017.

